# Structural basis of RECQL5-induced RNA polymerase II transcription braking and subsequent reactivation

**DOI:** 10.1101/2025.01.29.635449

**Authors:** Luojia Zhang, Yuliya Gordiyenko, Tomos Morgan, Catarina Franco, Ana Tufegdžić Vidaković, Suyang Zhang

**Affiliations:** MRC Laboratory of Molecular Biology; Cambridge, CB2 0QH, United Kingdom

**Keywords:** Transcription, DNA repair, cryo-EM, elongation rate

## Abstract

During productive transcription elongation, the speed of RNA polymerase II (Pol II) is highly dynamic within individual genes and varies between different genes^1,2^. Unregulated rapid transcription elongation can lead to detrimental consequences such as transcription-replication collisions, altered alternative splicing patterns, and genome instability^1–7^. Therefore, elongating Pol II requires mechanisms to slow its progression, yet the molecular basis of transcription braking remains unclear. RECQL5 is a DNA helicase that functions as a general elongation factor by slowing down Pol II^8–11^. Here we report cryo-electron microscopy (cryo-EM) structures of human RECQL5 bound to multiple transcription elongation complexes. Combined with biochemical analysis, we identify an α-helix of RECQL5 responsible for Pol II binding and slowdown of transcription elongation. We further reveal that the transcription-coupled DNA repair (TCR) complex allows Pol II to overcome RECQL5-induced transcription braking through concerted actions of its translocase activity and competition with RECQL5 for engaging Pol II. Additionally, RECQL5 inhibits TCR-mediated Pol II ubiquitination to prevent activation of the DNA repair pathway. Our results suggest a model in which RECQL5 and the TCR complex coordinately regulate the transcription elongation rate to ensure transcription efficiency while maintaining genome stability. This work provides a framework for future studies on the regulatory role of elongation speed in gene expression.

## Introduction

During eukaryotic transcription, release from the promotor-proximal pausing region is mediated by the positive transcription elongation factor-b (P-TEFb) that allows the formation of an activated elongation complex (EC*) to commence productive elongation^12,13^. EC* allows Pol II to drastically increase its speed and processivity, and contains additionally the elongation factors DSIF (a two subunit complex of SPT4 and SPT5), SPT6 and PAF (composed of CTR9, PAF1, SKI8, LEO1 and CDC73)^13^. However, the Pol II elongation rate is highly dynamic in cells and varies within individual genes and between different genes^1,2^. Transcription elongation rate affects co-transcriptional processes such as splicing, termination, mRNA modification and genome stability^1–7,14,15^. While a slow elongation rate was observed over exons, exon-intron junctions and poly(A) sites^14–18^, the mechanism underlying this transcription braking during productive elongation remains unclear.

RECQL5 was identified as a general elongation factor that controls transcription elongation rate^10^. Previous studies showed that RECQL5 decreases the Pol II elongation speed in vivo, inhibits transcription initiation and elongation in vitro and is crucial for maintaining genome stability^8–11,19^. Unlike other members of the RecQ helicase family, RECQL5 is only present in higher eukaryotes and uniquely interacts with Pol II^11,20^. Mice deficient in RECQL5 have increased risks of developing cancers, with profound sister chromatid exchange and double stranded DNA breaks^21,22^. Additionally, RECQL5 plays a critical role at the intersection of transcription, replication and DNA recombination^22–28^.

Here we report cryo-EM structures of human RECQL5 bound to multiple transcription elongation complexes. Together with biochemical analysis, we define the molecular basis of RECQL5-induced transcription slowdown. Furthermore, we identify that the transcription-coupled DNA repair (TCR) complex reactivates Pol II following RECQL5- induced transcription braking and restores the elongation rate. Our results provide a mechanistic understanding of the interplay between RECQL5 and the TCR complex in regulating Pol II elongation rate, thereby maintaining genome stability.

## Results

### Structure of the RECQL5-transcription complexes

All RecQ helicases contain a core helicase domain followed by the RecQ C-terminal (RQC) domain. RECQL5 has a unique C-terminal extension that consists of two protein-interacting domains, an internal Pol II-interacting (IRI) domain and a Set2-Rpb1-interacting (SRI) domain (Fig. 1a). The IRI domain contains the kinase-inducible domain interacting (KIX) domain. While the SRI domain was suggested to interact with the flexible phosphorylated Pol II C-terminal domain (CTD)^29–31^, a medium-resolution cryo-EM reconstruction revealed that the KIX domain may bind to the Pol II jaw^9^.

**Fig 1.**
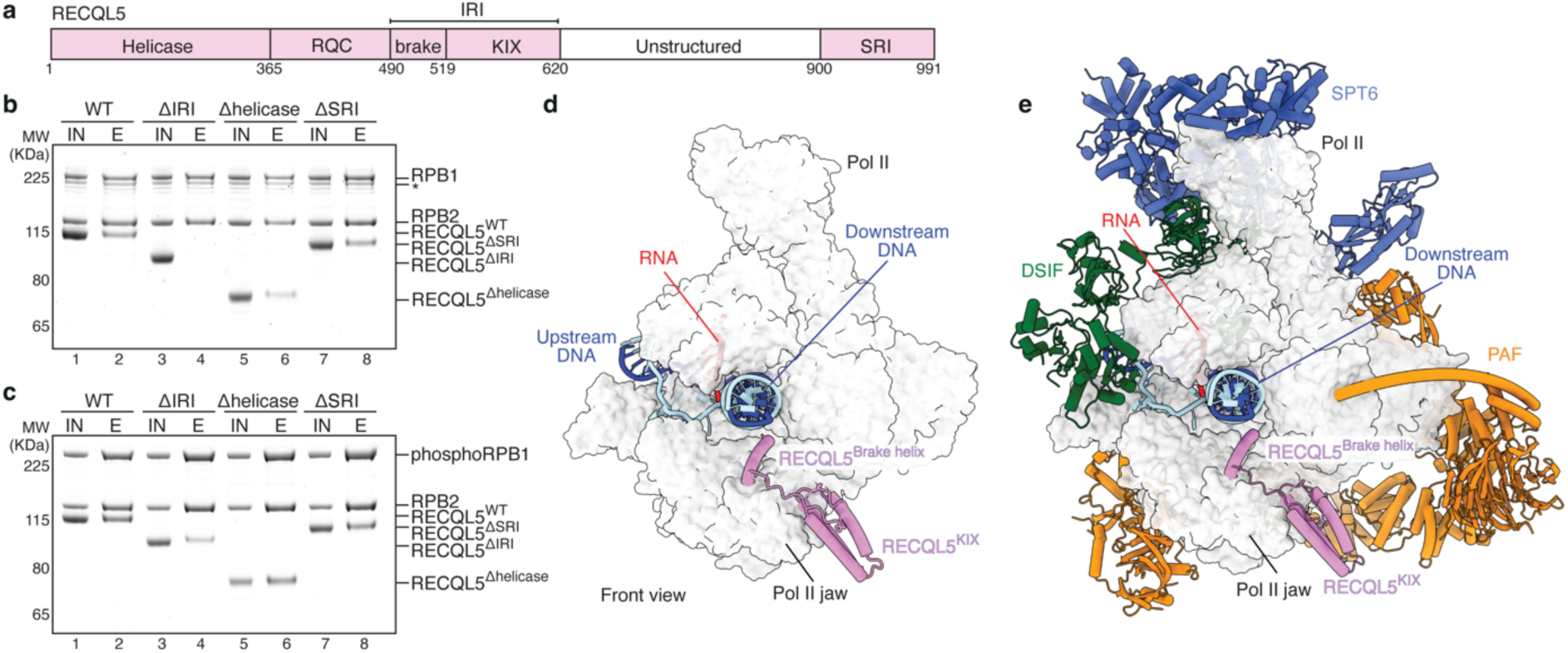
Structures of the EC-RECQL5 and EC*-RECQL5 complexes. **a**, Domain organization of RECQL5. brake: brake helix. **b**, Pulldown of non-phosphorylated Pol II with different RECQL5 constructs using the Twin-Strep tag on Pol II. Asterisk indicates RPB1 with CTD degradations. IN: input, E: elution. **c**, Pulldown of phosphorylated Pol II with different RECQL5 constructs. All pulldowns were repeated in triplicate. **d**, Structure of the EC-RECQL5 complex in front view. Pol II is shown in light grey surface representation, with RECQL5 shown in pink cartoon. Template DNA is coloured dark blue, non-template DNA in cyan and RNA in red. **e**, Structure of the EC*-RECQL5 complex with the elongation factors SPT6 (blue), DSIF (green) and PAF (orange) in cartoon representation.

To gain insights into the interactions between Pol II and RECQL5, we performed pulldowns using a Twin-Strep-tagged human Pol II and RECQL5 constructs with individual domains (helicase, IRI or SRI) deleted (Fig. 1a-c). In the absence of Pol II phosphorylation, deletion of the IRI domain eliminated RECQL5 binding to Pol II (Fig. 1b lane 4), indicating that the IRI domain engages the Pol II body and does not require Pol II phosphorylation for binding, consistent with previous studies^8,9,32^. Deletion of either the helicase domain or the SRI domain had no effect on RECQL5 binding to non-phosphorylated Pol II.

During transcription elongation, the Pol II CTD is hyperphosphorylated^33,34^. Compared to non-phosphorylated Pol II, we observed a slight increase in RECQL5 binding to in vitro phosphorylated Pol II (Fig. 1c lane 2). This increase is contributed by the interaction between the RECQL5 SRI domain and the phosphorylated Pol II CTD (Extended Data Fig. 1a), which enhances the affinity of RECQL5 to Pol II through multiple interaction sites. Deletion of the IRI domain strongly reduced RECQL5 binding to phosphorylated Pol II, whereas loss of the SRI domain resulted in a minor decrease of bound RECQL5 (Fig. 1c compare lane 4 to 8). Overall, these results demonstrate that the IRI domain confers the major contribution for RECQL5 interactions with Pol II, whereas the SRI domain enhances binding to phosphorylated Pol II.

To define the molecular basis of the interactions between RECQL5 and Pol II, we determined cryo-EM structures of RECQL5 bound to both a transcription elongating Pol II (EC) and the activated transcription elongation complex EC* (Fig. 1d, e, Supplementary Video 1). To reconstitute the EC-RECQL5 and EC*-RECQL5 complexes, recombinantly purified human RECQL5 was incubated with pre-assembled phosphorylated transcription elongation complexes and subjected to single particle cryo-EM analysis (Extended Data Figs. 1-5, Extended Data Table 1). The cryo-EM reconstructions reached an overall resolution of 2.8 Å for the EC-RECQL5 complex and 2.0 Å for the EC*-RECQL5 complex. We observed cryo-EM densities corresponding to the RECQL5 IRI domain in both reconstructions with a local resolution around 3 Å. Modelling of the EC and EC* structures^13,35^ and fitting of AlphaFold predictions of RECQL5 and elongation factors resulted in models of the EC-RECQL5 and EC*-RECQL5 complexes with good stereochemistry (Fig. 1d, e, Extended Data Table 1).

### The RECQL5 brake helix is crucial for Pol II binding

The structures reveal that RECQL5 binds directly to the jaw domain of the Pol II RPB1 subunit via its KIX domain (Fig. 1d, e, Fig. 2a-c). Additionally, an ⍺-helix (residues 490-519) N-terminal to the KIX domain rests on the jaw domain of RPB1 and extends towards the minor groove of the downstream DNA. Basic residues at the N-terminus of the helix, R496 and R502, are located near the DNA phosphate backbone (Extended Data Fig. 5b). We named this ⍺-helix of RECQL5 the brake helix. Therefore, the IRI domain of RECQL5 comprises the brake helix and the KIX domain (Fig. 1a). Strikingly, deleting the brake helix completely abolished RECQL5 binding to unphosphorylated Pol II, underlining the importance of the brake helix in the Pol II-RECQL5 interaction (Fig. 2d).

**Fig 2.**
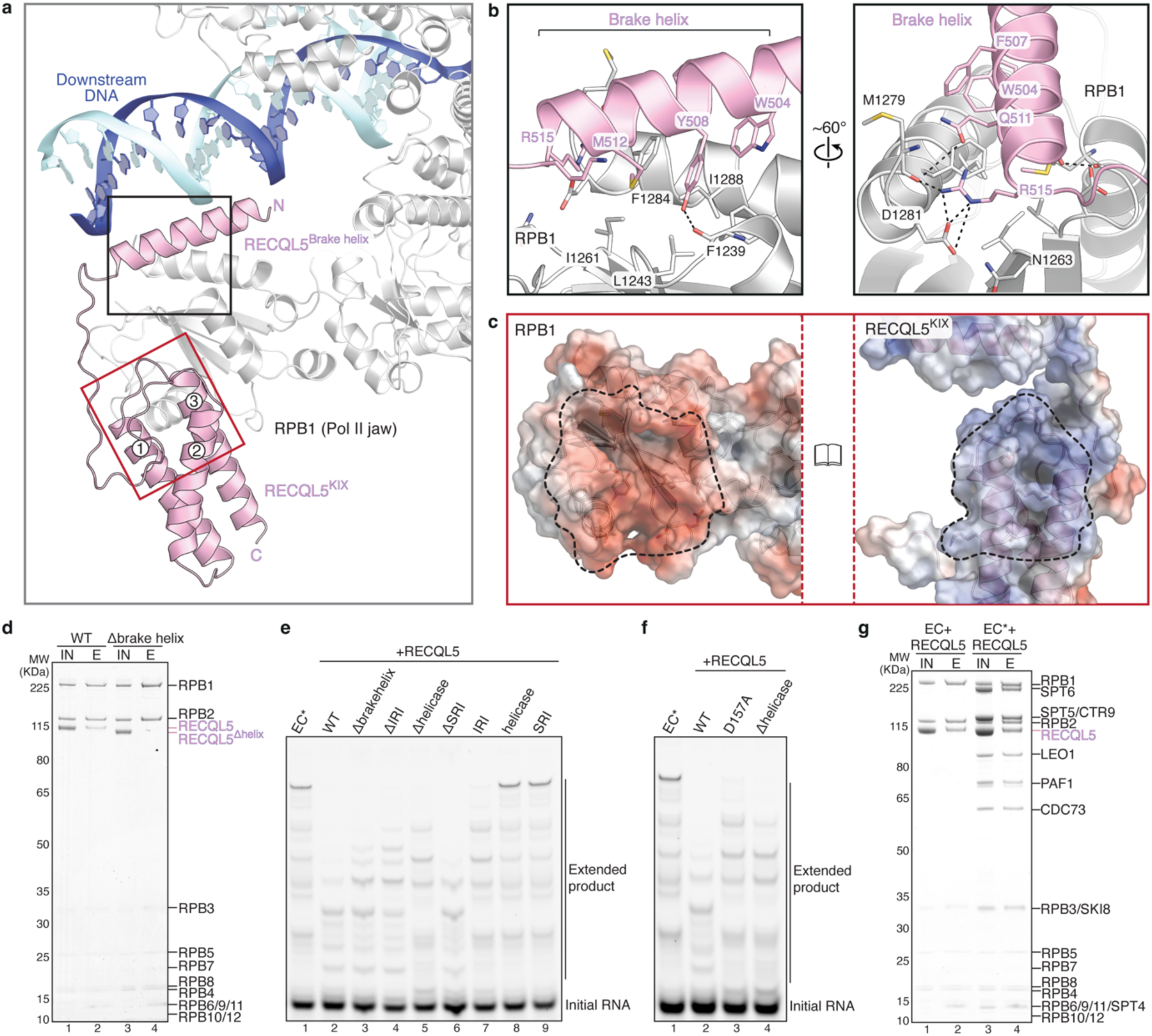
The RECQL5 brake helix is crucial for Pol II binding and transcription slowdown. **a**, Close-up view of interactions between RECQL5 (pink) and the jaw domain of the Pol II RPB1 subunit (light grey). The brake helix protrudes towards the downstream DNA (blue and cyan). **b**, Detailed views of interactions between the RECQL5 brake helix and the RPB1 jaw domain. **c**, The electrostatic surface potential of the RPB1 jaw domain (left) and the RECQL5 KIX domain (right) in book view with a range of ± 5 kT/e, where blue and red represent positively and negatively charged areas, respectively. The dashed lines highlight the interface areas between RPB1 and RECQL5 that are oppositely charged. **d**, Pulldown of non-phosphorylated Pol II with wildtype and brake-helix-deleted RECQL5. IN: input, E: elution. **e**, **f**, RNA extension assays of EC* in the presence of various RECQL5 constructs showing that the brake helix and the helicase activity of RECQL5 are critical for slowing down transcription elongation. **g**, Pulldown of EC and EC* with RECQL5 using the Twin-Strep tag on Pol II. All assays and pulldowns were repeated in triplicate.

The brake helix of RECQL5 forms extensive contacts with the jaw domain of RPB1 (Fig. 2a, b). Bulky side chains of W504, F507, Y508 and M512 on the brake helix engage a hydrophobic surface of RPB1 that is formed by F1239, L1243, I1261, F1284 and I1288 (Fig. 2b, Extended Data Fig. 5a). The brake helix interaction with the RPB1 jaw is further stabilized by multiple hydrogen bonds, including R515 of the brake helix with D1281 and M1279 of RPB1, Q511 of the brake helix with RPB1 M1279, and Y508 of the brake helix with RPB1 F1239 (Fig. 2b, Extended Data Fig. 5a).

The interface between the RECQL5 KIX domain and RPB1 shows charge complementarity (Fig. 2c). A negatively charged surface of RPB1 formed by glutamates and aspartates docks onto a positively charged patch on helices 1 and 3 of the KIX domain (Fig. 2a and c).

Residues at the interface of RECQL5 and RPB1 are highly conserved among higher eukaryotes (Extended Data Fig. 6), indicating that RECQL5 may likewise interact with Pol II in other eukaryotic species. Crosslinking-coupled mass spectrometry data identified crosslinks between the RECQL5 IRI domain and Pol II jaw, further validating the structural data (Extended Data Fig. 7a-c, Extended Data Table 2).

### RECQL5 induces transcription braking

Previous studies reported that RECQL5 inhibits transcription initiation and elongation in vitro in the presence of initiation factors^8,9^. Both the IRI and helicase domains were found to be important for the inhibitory effect of RECQL5, but the helicase activity was not required^8^. To understand the role of RECQL5 in transcription elongation, we recapitulated productive transcription elongation using the EC* complex containing elongation factors DSIF, SPT6 and PAF (Fig. 2e).

RECQL5 slows down transcription elongation effectively, resulting in accumulation of shorter transcripts and Pol II stalling at different locations compared to EC* transcription in the absence of RECQL5 (Fig. 2e, compare lane 2 to 1). Deletion of either the brake helix alone or the entire IRI domain partially relieved the RECQL5-induced transcription braking (Fig. 2e, lanes 3 and 4), indicating that the brake helix plays a critical role in slowing down Pol II. The brake helix extends towards the minor groove of the downstream DNA, possibly providing a steric hindrance for Pol II to proceed with transcription (Fig. 2a, Extended Data Fig. 5b). Interestingly, deletion of the helicase domain not only partially restored transcription but also recovered the Pol II stalling pattern to that of EC* (Fig. 2e, compare lane 5 to 1). On the other hand, although Pol II is phosphorylated in EC*, loss of the SRI domain did not impact the inhibitory effect of RECQL5 on transcription (Fig. 2e lane 6). The IRI domain alone moderately slowed down transcription elongation, whereas neither the helicase domain nor the SRI domain in isolation affected transcription (Fig. 2e, lanes 7-9).

As we could not observe cryo-EM density corresponding to the RECQL5 helicase domain, it may induce transcription braking either as a roadblock for Pol II movement by binding to the downstream DNA or as a helicase to partially unwind the downstream DNA. To investigate whether the helicase activity of RECQL5 is required for its inhibitory effect on productive transcription elongation, we used RECQL5^D157A^, a previously characterized helicase mutant^8^. Electromobility shift assay (EMSA) showed that the D157A mutation does not affect RECQL5 binding to DNA (Extended Data Fig. 8a). Strikingly, the RECQL5^D157A^ mutant failed to slow down transcription to the same extent as wildtype RECQL5 (Fig. 2f).

Instead, both the Pol II stalling pattern and the level of transcription braking by RECQL5^D157A^ resemble that of RECQL5^Δhelicase^ (Fig. 2f, compare lanes 3 and 4). These data reveal that the helicase activity of RECQL5 is important for slowing down productive transcription elongation, which differs from previous transcription assays using initiation factors. The RECQL5 helicase activity may contribute to the change in Pol II stalling pattern by partially unwinding and distorting the downstream DNA. Overall, our results show that, while the brake helix is essential for RECQL5 binding to Pol II, both the brake helix and the helicase activity of RECQL5 are required to slow down transcription during productive elongation.

### RECQL5 binding accommodates elongation factors

Our structures reveal that RECQL5 engages both EC and EC* identically on the Pol II jaw via its IRI domain and does not change the binding modes of elongation factors DSIF, SPT6 and PAF on Pol II, which are required for productive elongation (Fig. 1d, e).

Additionally, we find that the presence of elongation factors does not impair RECQL5 binding to EC (Fig. 2g). Furthermore, RECQL5 forms a complex with the elongation factor SPT6 in our pulldown assay and interacts weakly with DSIF and PAF (Extended Data Fig. 7d). Supporting this, our crosslinking-coupled mass spectrometry data showed interlinks of RECQL5 with elongation factors SPT6, DSIF and PAF (Extended Data Fig. 7a-c, Extended Data Table 2). These results reveal that RECQL5 binding is compatible with EC* and the elongation factors may be involved in the recruitment of RECQL5 to Pol II.

Although RECQL5 causes transcription elongation to proceed at a lower speed, it does not arrest Pol II to a complete stop. The active site of Pol II adopts a post-translocated state that can accept incoming nucleotide substrate (Extended Data Fig. 5g). In addition, superposition of the EC*-RECQL5 structure with a structure of EC* transcribing into a nucleosome^36^ revealed that RECQL5 binding at the Pol II jaw has no clash with the downstream nucleosome (Extended Data Fig. 8b).

### RECQL5 and UVSSA have overlapping binding sites on Pol II

The binding site of the RECQL5 KIX domain on the Pol II jaw overlaps with that of transcription factor IIS (TFIIS)^37^ (Extended Data Fig. 8c) and UV-stimulated scaffold protein A (UVSSA)^38^, a component of the transcription-coupled DNA repair (TCR) complex (Fig. 3a, b, Extended Data Fig. 9a). TFIIS helps Pol II to overcome stalling by stimulating cleavage of backtracked RNA, allowing reactivation of transcription elongation^37^. Previous studies found that RECQL5 inhibited TFIIS-mediated transcription readthrough of elongation blocks^9^ and UVSSA prevented TFIIS-mediated cleavage of backtracked RNA^38^. Nevertheless, the functional relationship between RECQL5 and UVSSA remains unclear.

**Fig 3.**
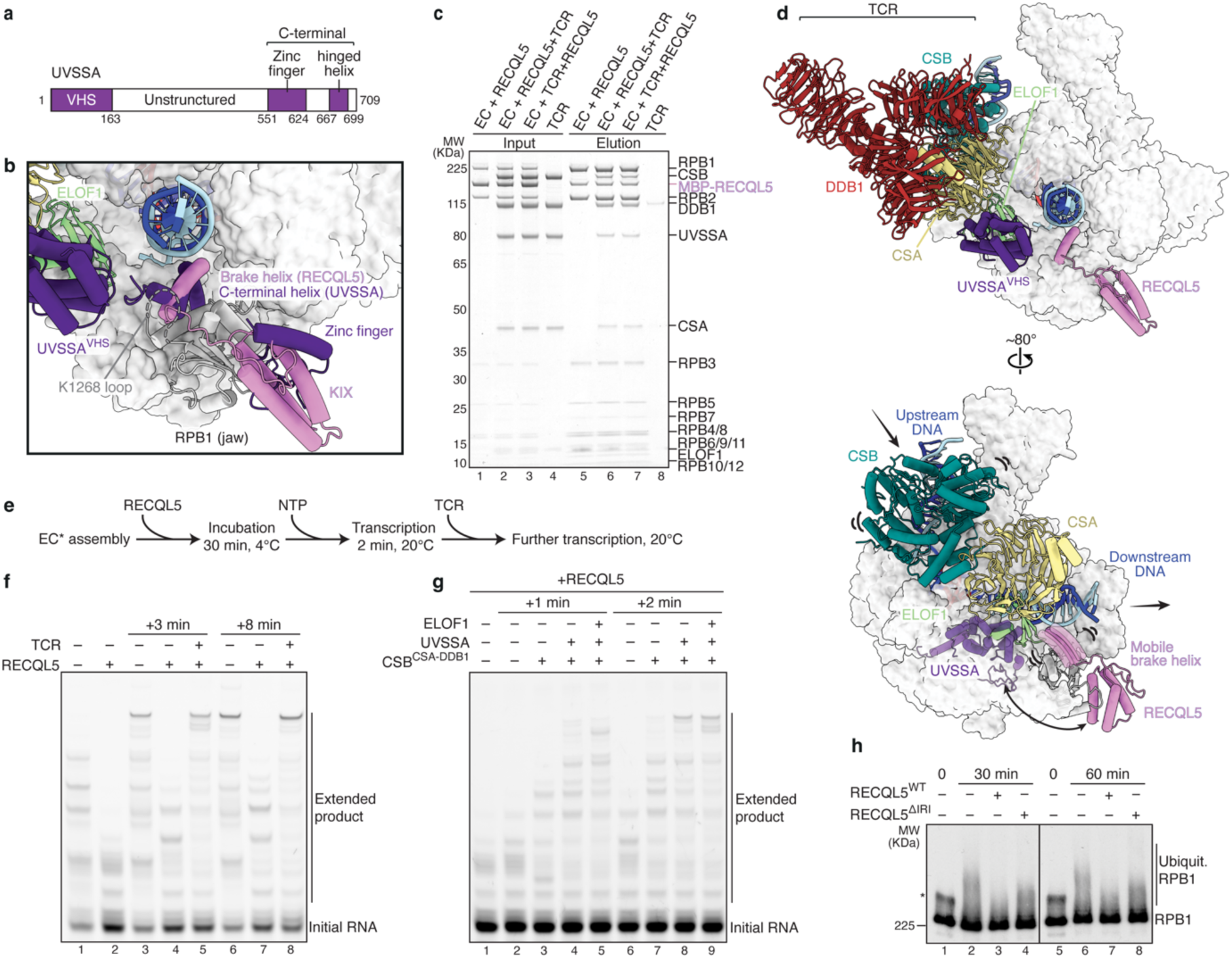
The TCR complex reactivates transcription elongation. **a**, Domain organization of UVSSA. **b**, Superposition of the EC-RECQL5 structure with the EC-TCR structure (PDB: 8B3D)^38^ revealed that the RECQL5 brake helix (pink) and the C-terminal hinged helix of UVSSA (dark purple) bind to the same site on the Pol II jaw (grey), while the RECQL5 KIX domain and the UVSSA zinc finger domain occupy the same binding site. The two helices are positioned next to the K1268 loop which is the main target of Pol II ubiquitination. **c**, RECQL5 and the TCR complex can bind to Pol II simultaneously. The competition assay was performed using the Twin-Strep tag on Pol II, with either RECQL5 or the TCR complex added to Pol II first. **d**, Cryo-EM structure of the EC-TCR-RECQL5 complex. Pol II is shown in light grey surface representation, while RECQL5 (pink) and the TCR components CSB (teal), CSA (yellow), DDB1 (dark red), ELOF1 (lime), UVSSA (dark purple) are shown as cartoon. Lower panel: The CSB translocase activity combined with competition by UVSSA makes the RECQL5 brake helix mobile. UVSSA is depicted in transparent cartoon at its approximate location to illustrate the potential competition with RECQL5. DDB1 is hidden to provide a clear view of the brake helix. **e**, Schematic of the EC* RNA extension assay in the presence of RECQL5 and the TCR complex. **f**, The TCR complex reactivates elongation following RECQL5-induced transcription braking. Initial transcription proceeded for 2 minutes without or with RECQL5 (lane 1 and 2) before addition of the TCR complex and further transcription for 3 or 8 minutes. **g**, Both CSB and UVSSA are important to restore transcription elongation. Initial transcription with RECQL5 proceeded for 2 minutes before addition of TCR factors and further transcription for 1 or 2 minutes. **h**, RECQL5 inhibits TCR-mediated Pol II ubiquitination shown on the Western blot using an antibody against RPB1. Asterisk indicates endogenously phosphorylated RPB1 at 0 timepoint. All assays were repeated in triplicate.

Besides UVSSA, the TCR complex contains the Cockayne syndrome proteins, CSB and CSA, and the DNA damage binding protein 1 (DDB1). Transcription elongation factor 1 homolog (ELOF1) helps to stably position UVSSA on the Pol II jaw, allowing targeted Pol II ubiquitination and activation of the DNA repair pathway^38,39^. The TCR complex interacts with Pol II via three contact points mediated by CSB, UVSSA and ELOF1 (Extended Data Fig. 9a)^38^. CSB competes with DSIF for the binding site on the Pol II stalk, thereby displacing DSIF from Pol II^38^.

A comparison of our EC-RECQL5 structure and the EC-TCR structure^38^ revealed that not only the RECQL5 KIX domain and the UVSSA zinc finger domain bind to the same site on the Pol II jaw, but the RECQL5 brake helix also aligns perfectly onto the C-terminal hinged helix of UVSSA, extending into the minor groove of the downstream DNA (Fig. 3a, b, Extended Data Fig. 9a). The RECQL5 brake helix and the UVSSA C-terminal helix have opposite directionality (Extended Data Fig. 9a right panel). While both helices use an arginine (UVSSA: R669 and RECQL5: R515) at the end of the helix to form hydrogen bonds with the Pol II jaw, RECQL5 engages the hydrophobic patch of the Pol II jaw with bulky aromatic residues, which are replaced by an isoleucine and a valine in UVSSA. The overlapping binding sites of RECQL5 and UVSSA on Pol II, combined with the ability of CSB to promote Pol II forward translocation on non-damaged templates^40–42^, prompted us to investigate the role of the TCR complex in restoring RECQL5-induced transcription slowdown.

### Structure of the EC-TCR-RECQL5 complex

We first tested whether the TCR complex can displace RECQL5 from Pol II using competing assays. The competition assays were performed in the absence of phosphorylation to eliminate the SRI-CTD interaction, thereby restricting RECQL5 association with Pol II to the IRI-jaw interaction, which overlaps with the UVSSA binding site. UVSSA only interacted weakly with Pol II in the presence of ELOF1 (Extended Data Fig. 9b). Addition of RECQL5^ΔSRI^ to the pre-assembled EC-UVSSA complex showed that more RECQL5 than UVSSA was bound to EC, indicating that the RECQL5 IRI domain is a stronger binder to the Pol II jaw compared to the UVSSA C-terminal region. We further tested the ability of the complete TCR complex to displace RECQL5, in which either RECQL5 or the TCR complex was pre-incubated with EC to form a complex before addition of the competitor (Fig. 3c).

The amount of RECQL5 and TCR factors bound to Pol II remained the same across all conditions, regardless of the order in which RECQL5 and the TCR complex were added. Despite occupying the same binding site on the Pol II jaw, RECQL5 binding to Pol II is not diminished by the TCR complex, likely due to the IRI domain being a stronger binder. Both the TCR complex and RECQL5 can co-exist on the Pol II surface as the TCR complex engages Pol II additionally via CSB and ELOF1 (Extended Data Fig. 9a).

To further investigate whether the TCR complex and RECQL5 can bind to Pol II simultaneously, we determined the cryo-EM structure of an EC-TCR-RECQL5 complex with an overall resolution of 3.5 Å (Fig. 3d, Extended Data Fig. 10, Supplementary Video 1, Extended Data Table 1). The TCR complex engages Pol II in the same mode as the EC-TCR structure^38^, with CSB interacting with the upstream DNA and Pol II clamp, while ELOF1 binds stably to the RPB2 lobe near the DNA entry tunnel. RECQL5 binds to the Pol II jaw identically as in the EC-RECQL5 structure, with the brake helix extending towards the downstream DNA. On the other hand, only weak cryo-EM densities corresponding to the UVSSA N-terminal VHS domain were observed (Extended Data Fig. 10). The structure reveals that both the TCR complex and RECQL5 can bind to Pol II at the same time, although RECQL5 prevents the stable engagement of the UVSSA C-terminal region to the Pol II jaw.

### The TCR complex reactivates transcription elongation

We next investigated whether the TCR complex can restore transcription elongation following RECQL5-induced transcription braking in vitro. In this assay, we allowed RECQL5-mediated transcription slowdown to progress for two minutes before addition of the TCR components to proceed with further transcription (Fig. 3e-g). In the absence of TCR factors, elongation is slow due to transcription braking by RECQL5. Addition of CSB-CSA- DDB1 partially relieved the transcription slowdown, whereas addition of the entire TCR complex including UVSSA and ELOF1 completely restored transcription to the same level as EC* in the absence of RECQL5 (Fig. 3f, g). Re-activation of transcription elongation is mediated by the translocase activity of CSB, which binds to the upstream DNA and pushes Pol II towards the downstream DNA^40,43^. Additionally, the UVSSA C-terminal hinged helix, stabilized by ELOF1, competes for the same binding site as the RECQL5 brake helix on the Pol II jaw. Despite being a weaker binder, the concerted actions of CSB and UVSSA may distort or displace the RECQL5 brake helix, thereby relieving the RECQL5-induced transcription slowdown (Fig. 3d).

### RECQL5 inhibits TCR-mediated Pol II ubiquitination

When Pol II encounters DNA damage, recruitment of the TCR complex triggers Pol II ubiquitination and activation of the DNA repair pathway^44,45^. K1268 located on a loop of the jaw domain of RPB1 subunit is the main target for Pol II ubiquitination, serving as a master switch to regulate transcription and DNA repair^46,47^. Correct positioning of the UVSSA C-terminal hinged helix next to the K1268 loop is required for accurate Pol II ubiquitination^38^. Our structures reveal that the brake helix of RECQL5 competes with the UVSSA C-terminal hinged helix for the same binding site next to the K1268 loop (Fig. 3b). We therefore hypothesized that, while the TCR complex restores the transcription elongation speed, RECQL5 may prevent Pol II ubiquitination to avoid excessive triggering the DNA repair pathway when no DNA damage is detected.

To investigate the effect of RECQL5 on TCR-mediated Pol II ubiquitination, we performed in vitro Pol II ubiquitination assays using the E3 ligase CRL4 (Fig. 3h, Extended Data Fig. 9c). The E3 ligase CRL4 targets both Pol II and CSB for ubiquitination in response to UV damage, triggering the DNA repair pathway^46–48^. Remarkably, RECQL5 specifically inhibits RPB1 ubiquitination without affecting CSB ubiquitination, demonstrating the additional role of RECQL5 in preventing excessive Pol II ubiquitination during transcription elongation in the absence of DNA damage. Deleting the IRI domain strongly reduced the inhibitory effect of RECQL5 (Fig. 3h), supporting the idea that RECQL5 prevents Pol II ubiquitination by competing with UVSSA for the same binding site on the Pol II jaw.

## Discussion

It is increasingly evident that transcription elongation is far more complex than initially thought, with transcription elongation serving as a key regulatory step in gene expression.

Uncontrolled Pol II elongation speed disrupts co-transcriptional processes and compromises genome stability, with mutations affecting Pol II speed found to be embryonic lethal in mice^1–7,14,15,49^. Furthermore, transcription speed has been shown to adapt in response to internal and external stimuli^50–52^. Nevertheless, the mechanism by which Pol II decelerates and re- accelerates during productive elongation remains poorly understood.

In this study, we define the molecular basis of transcription braking by the elongation factor RECQL5 and reactivation of elongation by the transcription-coupled DNA repair complex (Fig. 4, Supplementary Video 1). Our structural and biochemical data suggest a model in which RECQL5 induces transcription braking by engaging the fast-moving EC* during productive elongation (Fig. 4a). RECQL5 slows down transcription via its brake helix that protrudes towards the downstream DNA and its helicase activity that may partially unwind the downstream DNA (Fig. 4b, c). Slow-moving Pol II triggers the recruitment of the TCR complex to probe for possible DNA damage (Fig. 4d). If no DNA damage is detected, the TCR complex reactivates transcription elongation through a joint effort of CSB and UVSSA. While CSB uses its translocase activity to push Pol II forward, UVSSA competes with RECQL5 for the same binding site on the Pol II jaw, resulting in distortion or displacement of the RECQL5 brake helix. At the same time, RECQL5 inhibits specifically TCR-mediated Pol II ubiquitination, but not that of CSB, thereby preventing excess activation of the DNA repair pathway (Fig. 4e). CSB polyubiquitination followed by degradation by the proteasome allows the release of the TCR complex from Pol II and the re-establishment of EC* (Fig. 4f)^48^. Although not critical in our in vitro experiments, the interaction of RECQL5 with the Pol II CTD greatly increases the local concentration of RECQL5 and may allow swift rebinding of RECQL5 to the Pol II body.

**Fig 4.**
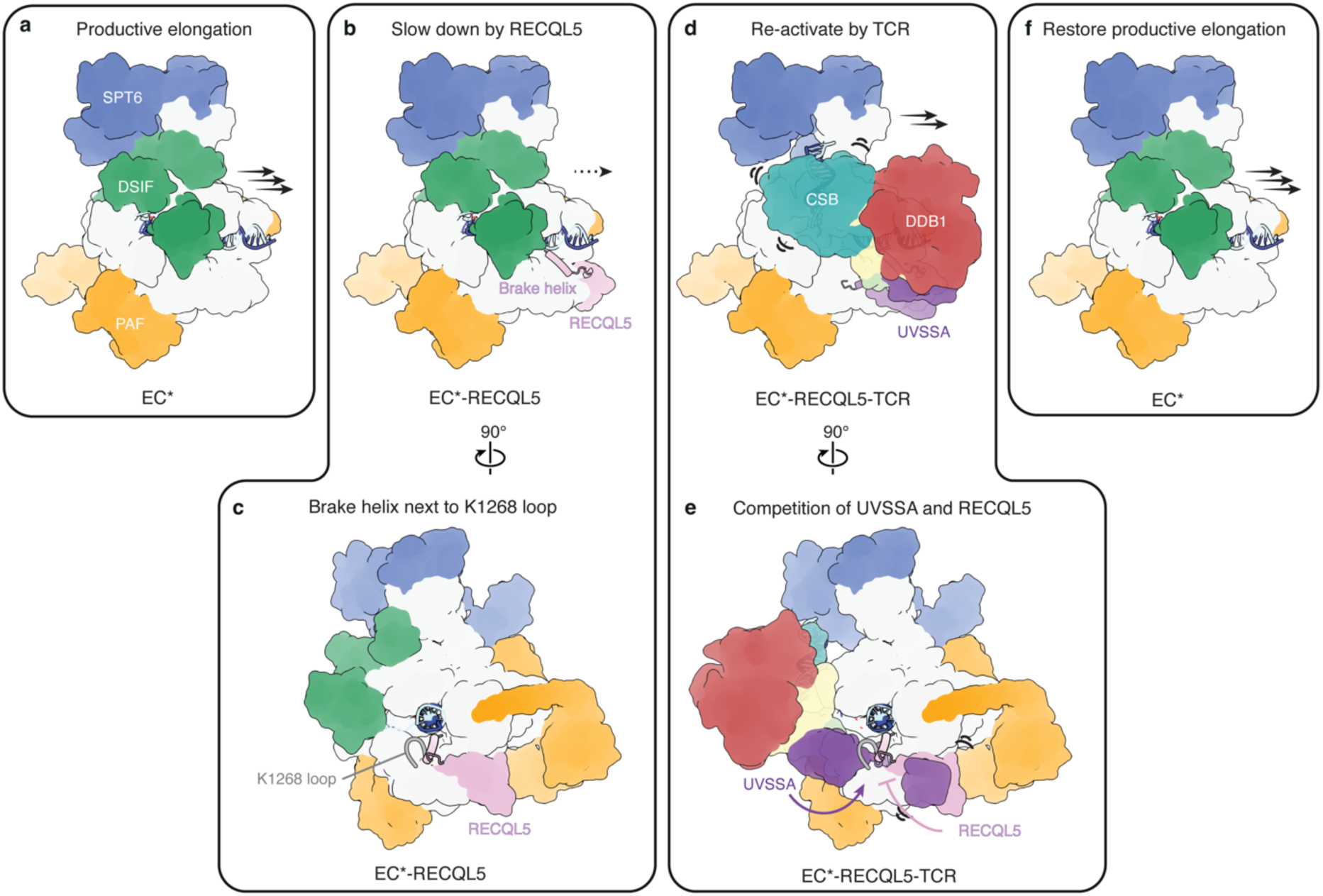
RECQL5 and the TCR complex coordinate to regulate the transcription elongation speed. RECQL5 recruitment to productive elongating EC* (**a**) slows down transcription through the brake helix and its helicase activity (**b**, **c**). Reactivation of transcription elongation requires the joint effort of CSB and UVSSA in the TCR complex, resulting in distortion of the RECQL5 brake helix (**d**). At the same time, RECQL5 inhibits TCR-mediated Pol II ubiquitination to prevent activation of the DNA repair pathway (**e**). CSB ubiquitination results in displacement of the TCR complex and restoration of EC* (**f**).

Supporting this model, loss of RECQL5 leads to increased transcription elongation rate and transcription-induced genome instability in vivo^10^, demonstrating the role of RECQL5 as a general elongation factor. In the absence of RECQL5, sites with elevated transcription elongation were enriched in double-stranded DNA breaks, which appears to be associated with replication^10,53,54^. Therefore, RECQL5 may slow down transcription elongation to avoid transcription-replication collisions. On the other hand, reduced transcription elongation was observed in Cockayne syndrome group B cells^55^ and CSB stimulated transcription elongation rate in vitro^40,41,56^, supporting a role of CSB in activating transcription elongation.

Furthermore, CSB was isolated from whole cell extracts as a part of a large complex that contains Pol II and was involved in regulating nucleosome positioning^42,57^.

Taken together, our data suggest that RECQL5 acts as a brake to slow down transcription elongation, while the TCR complex functions as an accelerator to overcome the transcription braking. The coordinated actions of RECQL5 and the TCR complex ensures efficient transcription elongation, while maintaining genome stability, and possibly preventing transcription-replication collision and allowing co-transcriptional events to occur. Our work provides a framework for future studies on regulation of transcription elongation rate and its regulatory roles in gene expression.

## Methods

### Cloning and protein expression

Human RECQL5, Pol II-CTD, CSA, CSB, UVSSA, DDB1, CUL4A, RBX1, ELOF1 and UbcH5b were amplified from human cDNA isolated from FreeStyle293 cells (Thermo Fisher). RECQL5, Pol II-CTD, CSB and CUL4A were cloned into the 438C vector (addgene no. 55220) with an N-terminal His_6_-MBP tag, CSA and RBX1 were cloned into the 438A vector with no tag (addgene no. 55218), DDB1 and UVSSA were cloned into the 438B vector (addgene no. 55219) with an N-terminal His_6_ tag, and ELOF1 and UbcH5b were cloned into the 1C vector (addgene no. 29654) with an N-terminal His_6_-MBP tag using ligation independent cloning^58^. CSA and DDB1 as well as CUL4A and RBX1 were further combined into single vectors.

Proteins cloned into the 438-vector series were expressed in High Five cells (Gibco). 1 L High Five cells in Sf-900^TM^ II SFM medium (Gibco) were infected with P2 virus and grown for 50 to 72 hours. Cells were harvested by centrifugation at 1000xg for 18 min and frozen in liquid nitrogen and stored at -80 °C before protein purification. All mutants of RECQL5 were generated using Quikchange^59^ and were expressed and purified as the wildtype proteins.

Proteins cloned into 1C vector were expressed in BL21 (DE3) RIL cells (Agilent) in LB medium. Expression was induced with 1 mM IPTG when the cell density reached an OD of 0.6-0.8. UbcH5b was expressed overnight at 18 °C, whereas ELOF1 was expressed at 37 °C for 4h before harvesting and storage at -80 °C.

### Protein purification

The porcine Pol II was purified from *Sus scrofa domesticus* thymus as described^35^. Human P-TEFb, PAF, DSIF, RTF1 and SPT6 were purified as described^13^, while human TCR proteins were purified as described^38^.

For all RECQL5 constructs, cell pellets were resuspended in lysis buffer (50 mM Tris-HCl pH 7.4, 500 mM NaCl, 30 mM Imidazole pH 8.0, 5% glycerol, 1 mM DTT) supplemented with 1 mM PMSF and EDTA-free protease inhibitor tablets (Roche). The resuspended cells were sonicated, clarified by centrifugation and loaded onto a 5 ml HisTrap HP column (Cytiva). The column was washed with lysis buffer and eluted with lysis buffer supplemented with 300 mM Imidazole. The eluate was applied onto a home-packed amylose column (NEB), washed with lysis buffer and eluted with amylose elution buffer (50 mM Tris-HCl pH 7.4, 300 mM NaCl, 50 mM maltose, 5% glycerol, 1 mM DTT). The His_6_-MBP tag was cleaved with TEV protease overnight and applied to a HisTrap HP column to remove TEV protease, undigested protein and His_6_-MBP tag. Flowthrough was collected and further purified with HiLoad 16/600 Superdex 200 pg or HiLoad 16/600 Superdex 75 pg column (Cytiva) in RECQL5 SEC Buffer (20 mM HEPES-NaOH pH 7.5, 200 mM NaCl and 1 mM DTT). The peak fractions were concentrated using Amicon Ultra Centrifugal Filters (Merck). For purification of His_6_-MBP tagged wildtype RECQL5, the eluate from amylose column was diluted four times with Buffer A (50 mM Tris-HCl pH 7.4, 5% glycerol, 1 mM DTT) and applied onto a HiTrap Heparin HP column (Cytiva). The protein was eluted with a gradient of Buffer A with 1 M NaCl before subjecting to a final size exclusion purification step on the HiLoad 16/600 Superdex 200 pg column (Cytiva) in RECQL5 SEC Buffer. Peak fractions were concentrated, frozen in liquid nitrogen and stored at -80 °C.

The His_6_-MBP-tagged human Pol II-CTD was purified the same way as RECQL5 with a HisTrap HP column (Cytiva) followed by the amylose column (NEB). Following amylose elution, protein was diluted to 100 mM NaCl with Buffer A, applied on a HiTrapQ HP column, and eluted with a gradient of Buffer A with 1 M NaCl. Peak fractions were concentrated, frozen in liquid nitrogen and stored at at -80 °C.

### Sample preparation for cryo-EM

EC-RECQL5 and EC*-RECQL5 complexes were formed on a DNA-RNA scaffold with a mismatch bubble of 11 nucleotides. The same RNA and template DNA were used for both complexes: RNA: 5’ - GAG AGG GAA CCC ACU - 3’. Template DNA: 5’ - GCT CCC AGC TCC CTG CTG GCT CCG AGT GGG TTC TGC CGC TCT CAA TGG - 3’. For EC-RECQL5, a non-template DNA of the same length as the template DNA was used: 5’ - CCA TTG AGA GCG GCC CTT GTG TTC AGG AGC CAG CAG GGA GCT GGG AGC - 3’.

For EC*-RECQL5, a non-template DNA with 15-nucleotide 3’ overhang was used: 5’ - CCA TTG AGA GCG GCC CTT GTG TTC AGG AGC CAG CAG GGA GCT GGG AGC CTT AGA CAG CAT GTC - 3’. All oligos were synthesized by IDT and resuspended in water.

For the EC-RECQL5 complex, RNA was annealed with equimolar amount of template DNA by incubating at 60 °C for 5 min, followed by a gradual decrease in temperature at a rate of 1 °C min^-1^ to a final temperature of 30 °C in 20 mM HEPES pH 7.5, 100 mM NaCl and 3 mM MgCl_2_. The porcine Pol II (75 pmol) was incubated with the RNA-DNA hybrid (150 pmol) at 30 °C for 10 min, followed by addition of the non-template DNA (300 pmol) and incubation at 30 °C for another 10 min. The complex was phosphorylated with GSK3B (300 pmol) and ATP (1 mM) in SEC100 Buffer (20 mM HEPES pH 7.5, 100 mM NaCl, 3 mM MgCl_2_, 1 mM DTT) at 30 °C for 30 min, followed by incubation with RECQL5 (300 pmol) on ice for 30 min. The assembled EC-RECQL5 complex was applied to a Superdex 200 Increase 3.2/300 column (Cytiva) equilibrated in SEC150 Buffer (20 mM HEPES pH 7.5, 150 mM NaCl, 3 mM MgCl_2_ and 1 mM DTT). The peak fraction of EC- RECQL5 was used for freezing grids.

For the EC*-RECQL5 complex, the Pol II elongation complex was prepared essentially as described for the EC-RECQL5 complex. Final protein amounts used for complex formation were 100 pmol porcine Pol II, 200 pmol RNA-DNA hybrid and 400 pmol non-template DNA. The complex was mixed with elongation factors (SPT6, DSIF and PAF, 200 pmol each) and phosphorylated with P-TEFb (33 pmol) and ATP (1 mM) in SEC100 Buffer for 30 min at 30 °C. The assembled EC* complex was applied to a Superose 6 Increase 3.2/300 column (Cytiva) equilibrated in K50 Buffer (20 mM HEPES pH 7.5, 50 mM KCl, 4 mM MgCl_2_, 1 mM DTT). The peak fraction of EC* was incubated with 2x molar excess of RECQL5 and 1 mM ADP on ice for 30 min, crosslinked with 0.05% (v/v) glutaraldehyde for 45 min on ice and used directly for grid freezing.

The EC-TCR-RECQL5 complex was formed on a DNA-RNA scaffold with a mismatch bubble of 15 nucleotides^43^: Template DNA: 5’ - CGC TCT GCT CCT TCT CCC ATC CTC TCG ATG GCT ATG AGA TCA ACT AG - 3’. Non-template DNA: 5’ - CTA GTT GAT CTC ATA TTT CAT TCC TAC TCA GGA GAA GGA GCA GAG CG - 3’. RNA 5’ - ACA UCA UAA CAU UUG AAC AAG AAU AUA UAU ACA AAA UCG AGA GGA - 3’. The DNA and RNA oligos were synthesized by IDT. RNA was annealed with equimolar template DNA by incubating at 90 °C for 2 min, followed by a gradual decrease in temperature at a rate of 1 °C min^-1^ to a final temperature of 30 °C in 20 mM HEPES pH 7.5 and 100 mM NaCl. The porcine Pol II (75 pmol) was incubated with the RNA-DNA hybrid (150 pmol) at 30 °C for 10 min, followed by the addition of the non-template DNA (300 pmol) and incubation at 30 °C for an additional 10 min. The complex was mixed with elongation factors (SPT6 and PAF, 150 pmol each) and phosphorylated with P-TEFb (25 pmol) and ATP (1 mM) in SEC100 Buffer for 30 min at 30 °C before subjecting to a Superose 6 Increase 3.2/300 column (Cytiva) equilibrated in K50 Buffer. Peak fraction was incubated with 2x molar excess of RECQL5 on ice for 30 min, followed by incubation with 2x molar excess of TCR components (CSB, CSA-DDB1 and UVSSA), 4x molar excess of ELOF1 and 1 mM ADP at 30 °C for 10 min. The assembled EC-TCR-RECQL5 complex was crosslinked with 0.05% (v/v) glutaraldehyde for 45 min on ice and used directly for grid freezing.

2.5 μl of the sample was applied to the R3.5/1 carbon grids (Quantifoil) with a continuous carbon support (∼2.5 nm) for the EC-RECQL5 and EC*-RECQL5 complexes and no support layer for the EC-TCR-RECQL5 complex. The grids were glow-discharged for 13- 15 s by the Sputter Coater S150B (Edwards Vacuum). The grids were incubated with the sample for 30 s and blotted for 1-1.5s before plunge-freezing in liquid ethane with a Vitrobot Mark IV (Thermo Fisher) operated at 4 °C and 100% humidity.

### Pulldown assay

For pulldown of non-phosphorylated Pol II with different RECQL5 constructs, Twin-Strep-tagged human Pol II (12 pmol) was incubated with wildtype or mutant RECQL5 (36 pmol) on ice for 30 min in G-SEC150 Buffer (20 mM HEPES pH 7.5, 150 mM NaCl, 3 mM MgCl_2_, 5% glycerol, 1 mM DTT). Samples were incubated with the StrepTactinXT 4Flow high-capacity resin (IBA) at 4 °C for 2 hours. The resin was washed with G-SEC150 Buffer and eluted with G-SEC150 Buffer supplemented with 50 mM biotin. The eluate was separated on a 4-12% NuPAGE Bis-Tris gel (Invitrogen) and stained with InstantBlue (Abcam). Pulldowns using phosphorylated Pol II with different RECQL5 constructs were performed in the same way except that Pol II was incubated with P-TEFb (4 pmol) and ATP (1 mM) in SEC100 Buffer for 30 min at 30 °C before mixing with RECQL5 constructs in G- SEC150 Buffer.

For pulldown of Pol II-CTD with RECQL5^SRI^, His_6_-MBP-tagged Pol II-CTD (100 pmol) was phosphorylated with P-TEFb (25 pmol) and ATP (1 mM) in SEC100 Buffer for 30 min at 30 °C. P-TEFb storage buffer (20 mM HEPES pH 7.5, 300 mM NaCl, 10% glycerol, 1 mM DTT) is used instead of P-TEFb as a negative control for the non-phosphorylated CTD condition. The CTD was incubated with the SRI domain of RECQL5 (500 pmol) on ice for 30 min, followed by incubation with the amylose resin (NEB) equilibrated in G-SEC150 Buffer at 4 °C for 1 hour. The resin was washed with G-SEC150 Buffer and eluted with 20 mM maltose in G-SEC150 Buffer. The eluate was loaded on a 4-12% NuPAGE Bis-Tris gel (Invitrogen) and stained with InstantBlue (Abcam).

For pulldown of EC/EC* with RECQL5, the elongation complex was prepared in the same way as described for sample preparation for cryo-EM, except that Twin-Strep-tagged human Pol II was used. Final protein amounts used for complex formation were 12 pmol human Pol II, 24 pmol RNA-DNA hybrid and 48 pmol non-template DNA. The phosphorylation reaction was performed with P-TEFb (4 pmol) and ATP (1 mM) in SEC100 buffer for 30 min at 30 °C for either Pol II alone or with elongation factors (SPT6, DSIF and PAF, 24 pmol each). Samples were incubated with the StrepTactinXT 4Flow high-capacity resin (IBA) at 4 °C for 2 hours. The resin was washed with SEC100 Buffer and eluted with 50 mM biotin in SEC100 Buffer. The eluate was separated on a 4-12% NuPAGE Bis-Tris gel (Invitrogen), and stained with InstantBlue (Abcam).

For pulldown of RECQL5 with elongation factors SPT6, DSIF, PAF and RTF1, elongation factors (100 pmol) were phosphorylated with P-TEFb (17 pmol) and ATP (1 mM) in SEC100 Buffer for 30 min at 30 °C, followed by addition of His_6_-MBP-tagged RECQL5 (50 pmol) and incubation on ice for 30 min. Samples were incubated with the amylose resin (NEB) equilibrated in K50 Buffer at 4 °C for 1 hour. The resin was washed with K50 Buffer and eluted with K50 Buffer supplemented with 20 mM maltose. The proteins were separated on a 4-12% NuPAGE Bis-Tris gel (Invitrogen) and stained with InstantBlue (Abcam).

### RNA extension assay

All RNA extension assays were performed with a complementary DNA scaffold as follows: Template DNA: 5’ - CTG GAC TAC TGC GCC CTA GAC GTG CAG CAA GCT TGG GCT GCA GGT AAC CAG TTC TAC ATG CTA GAT ACT TAC CTG GTC GGA GGC CGA CGG - 3’. Non-template DNA: 5’ - CCG TCG GCC TCC GAC CAG GTA AGT ATC TAG CAT GTA GAA CTG GTT ACC TGC AGC CCA AGC TTG CTG CAC GTC TAG GGC GCA GTA GTC CAG - 3’. RNA: 5’ - /6-FAM/ - UUU UUU CCA GGU AAG -3’. All oligos were synthesized by IDT and resuspended in water.

RNA was annealed with equimolar amount of template DNA by incubating at 90 °C for 2 min, followed by a gradual decrease in temperature at a rate of 1 °C min^-1^ to a final temperature of 30 °C in 20 mM HEPES pH 7.5 and 100 mM NaCl. All concentrations refer to the final concentrations used in the assay. The porcine Pol II (150 nM) was incubated with the RNA-DNA hybrid (100 nM) at 30 °C for 10 min, followed by the addition of the non-template DNA (200 nM) and incubation at 30 °C for an additional 10 min. The complex was mixed with elongation factors (SPT6, DSIF and PAF, 150 nM each) and phosphorylated with P-TEFb (100 nM) and ATP (1 mM) in 20 mM HEPES pH 7.5, 100 mM NaCl, 3 mM MgCl_2_, 4% glycerol and 1 mM DTT at 30 °C for 30 min. The reactions were incubated with wildtype or mutant RECQL5 (1.5 µM) on ice for 30 min. Transcription extension was started by addition of 100 μM NTP at 20 °C. The reactions were quenched at various timepoints in equal volume of 2x stop buffer (6.4 M urea, 50 mM EDTA pH 8.0, 1x TBE). Reactions were treated with 0.13 unit/μl proteinase K (NEB) for 20 min at 37 °C before being applied onto 15% denaturing gels (15% acrylamide/bis-acrylamide 19:1, 7 M urea and 1x TBE, run in 0.5x TBE at 300 V for 100 min). The gels were visualized using the 6-FAM label on the RNA with the Typhoon 9500 FLA Imager (GE Healthcare). Extension assays with additional TCR factors were performed in the same way with the exception that EC* was incubated with 0.75 µM RECQL5, and 0.75 µM TCR factors was added to the reaction after extension by EC*-RECQL5 for 2 min.

### Electromobility shift assay

Electromobility shift assay for wildtype and D157A mutant RECQL5 was performed with two types of DNA scaffolds, one with a 3’ overhang and one with a splayed duplex.

DNAs were annealed in 20 mM HEPES pH 7.5 and 100 mM NaCl by heating at 94 °C for 4 min and slowly cool down to 30 °C at a rate of 1 °C min^-1^. Double-stranded DNAs at a final concentration of 0.5 μM were mixed with 0 to 2.5 μM of wildtype or D157A RECQL5 in 20 mM HEPES pH 7.5, 66 mM NaCl, 5% glycerol, 1 mM DTT, and 0.25 mg/ml BSA and incubated on ice for 30 min. The samples were then mixed with Novex Hi-Density TBE sample buffer (Thermo Fisher) and loaded onto a 6% DNA retardation gel (Invitrogen).

Electrophoresis was performed at 4 °C at 100 V in 0.5× TBE buffer for 105 min. Gels were stained with SYBR Gold (Thermo Fisher) and imaged with Typhoon 9500 FLA Imager (GE Healthcare).

### Pol II ubiquitination assay

Pol II elongation complexes were formed on the same DNA-RNA scaffold as for the cryo-EM studies of the EC-TCR-RECQL5 complex. RNA was annealed with equimolar template DNA by incubating at 60 °C for 5 min, followed by a gradual decrease in temperature at a rate of 1 °C min^-1^ to a final temperature of 30 °C in 20 mM HEPES pH 7.5, 100 mM NaCl and 3 mM MgCl_2_. All concentrations refer to the final concentrations used in the assay. Human Pol II (150 nM) was incubated with the RNA-DNA hybrid (300 nM) at 30 °C for 10 min, followed by addition of the non-template DNA (600 nM) and incubation at 30 °C for another 10 min. Subsequently, RECQL5 constructs (0 or 450 nM) was added to the elongation complex and incubated on ice for 30 min, before adding CSB (300 nM), CSA- DDB1 (300 nM), CUL4A-RBX1 (300 nM), ELOF1 (600 nM) and UVSSA (360 nM) for a further incubation at 30 °C for 15 min. Ubiquitination reactions were initiated upon addition of UBE1 (150 nM, R&D systems), UbcH5b (1.75 μM), ubiquitin (150 μM, R&D systems) and ATP (2 mM) in 50 mM Tris pH 8.0, 50 mM NaCl, 10 mM MgCl_2_ and 1 mM DTT at 37 °C. The reactions were quenched at various timepoints by mixing with 4x SDS loading dye (20 mM Tris-HCl pH 6.6, 8% SDS, 40% Glycerol, 0.8% bromophenol blue and 400 mM DTT). Reactions were loaded on a 3-8% Tris-acetate gel (Invitrogen) and transferred onto a 0.2 µm nitrocellulose membrane (Cytiva). The membrane was blocked with 5% (w/v) milk in PBS for 30 min at room temperature and incubated with F-12 anti-RPB1 antibody (1:1000 dilution, Santa Cruz Biotechnology, sc-55492) or anti-CSB antibody (1:1000 dilution, Santa Cruz Biotechnology, sc-166042) overnight at 4 °C. The membranes were washed with PBST (PBS with 0.2% Tween 20) and incubated with HRP-conjugated anti-mouse secondary antibody (1:10,000 dilution, Proteintech, SA00001-1) for 45 min at room temperature. The membranes were washed with PBST, developed with the Pierce ECL Chemiluminescent Substrate (Thermo Fisher) and filmed with OPTIMAX processor (Protec).

### Competition assay

Pol II elongation complex was prepared in the same way as described for the ubiquitination assays. Final protein amounts used for complex formation were 12 pmol human Pol II, 24 pmol RNA-DNA hybrid and 48 pmol non-template DNA. For the competition assay between RECQL5^ΔSRI^ and UVSSA-ELOF, the elongation complex was incubated with UVSSA-ELOF1 (60 pmol) at 30 °C for 15 min, followed by incubation with the StrepTactinXT 4Flow high-capacity resin (IBA) at 4 °C for 2 hours. The resin was washed with SEC100 Buffer to remove excess UVSSA-ELOF1 before addition of either RECQL5^ΔSRI^ (60 pmol) or RECQL5 SEC buffer, incubated at room temperature for 30 min, washed again with SEC100 Buffer, and eluted with SEC100 Buffer supplemented with 50 mM biotin. For the competition assay between RECQL5 and the TCR complex, the elongation complex was either incubated with RECQL5 (36 pmol) first on ice for 20 min, then with the TCR complex (36 pmol of CSB and CSA-DDB1, 50 pmol of UVSSA and ELOF1) on ice for 20 min in SEC100 Buffer, or vice versa. His_6_-MBP-tagged RECQL5 was used to avoid band overlapping with DDB1. The elongation complex was incubated with RECQL5 (36 pmol) alone in SEC100 Buffer on ice for 20 min as a positive control. The complexes were incubated with the StrepTactinXT 4Flow high-capacity resin (IBA) at 4 °C for 2 hours. The resin was washed with SEC100 Buffer with 5% glycerol and eluted with 50 mM biotin in SEC100 Buffer with 5% glycerol. The eluate was separated on a 4-12% NuPAGE Bis-Tris gel (Invitrogen) and stained with InstantBlue (Abcam).

### Cryo-EM data collection and processing

Cryo-EM data were collected on the 300 kV Titan Krios with a Falcon 4i direct electron detector (Thermo Fisher). Automated data acquisition was performed with EPU (Thermo Fisher) at a nominal magnification of 96,000x (0.8156 Å/pixel). Image stacks of 40 frames were collected with a defocus range of -0.5 μm to -2.0 μm in electron counting mode and a dose rate of 0.92-1.06 e^-^/Å^2^/frame. A total of 8,602 image stacks were collected for the EC-RECQL5 complex. A total of 60,808 image stacks were collected across two datasets for the EC*-RECQL5 complex. A total of 11,156 image stacks were collected for the EC-TCR- RECQL5 complex.

Motion correction and estimation of the contrast transfer function (CTF) were done in RELION 5.0 (refs ^60,61^). Particles in 400 pixels x 400 pixels for EC-RECQL5 and in 480 pixels x 480 pixels for EC*-RECQL5 and EC-TCR-RECQL5 were selected by automatic particle picking in Warp^62^. For EC-RECQL5 and EC*-RECQL5, further processing steps were performed in RELION 5.0 with final local refinement in CryoSPARC^63^. Further processing steps for EC-TCR-RECQL5 were performed in CryoSPARC.

For the EC-RECQL5 complex, two-dimensional (2D) classification, followed by three-dimensional (3D) refinement and local classification were performed to remove bad particles from the dataset (Extended Data Fig. 2a, b). Signal subtraction with a soft mask near the Pol II jaw (where the extra density of RECQL5 is) followed by focused 3D-classification without alignment was performed to separate EC-RECQL5 particles from EC-only particles (Extended Data Fig. 2c, d). Particles containing densities corresponding to RECQL5 were reverted to original particles and 3D-refined, followed by CTF refinement with per particle defocus estimation and Bayesian polishing to correct beam-induced particle motion (Extended Data Fig. 2e). Polished particles were subjected to an additional round of signal subtraction and focused 3D-classification on RECQL5 to further remove the EC-only particles (Extended Data Fig. 2f). Subsequent signal subtraction with a soft mask on the downstream DNA and focused 3D-classification without alignment were performed to exclude any apo Pol II-RECQL5 particles (Extended Data Fig. 2g, h). Particles containing densities of downstream DNA were reverted to original particles and imported into CryoSPARC for local refinement, resulting in the final EC-RECQL5 reconstruction with 66,771 particles at an overall resolution of 2.8 Å (Extended Data Fig. 2i).

For the two datasets of the EC*-RECQL5 complex, particles after cleanup in 2D and 3D classification were CTF-refined, and Bayesian-polished separately (Extended Data Fig. 3a-c, e-g). Polished particles in dataset 2 were further cleaned up by an additional round of local 3D-classification (Extended Data Fig. 3h, i). Signal subtraction with a soft mask on RECQL5 IRI domain and focused 3D-classification without alignment were performed to separate EC*-RECQL5 particles from EC*-only particles for both datasets (Extended Data Fig. 3d, j). Particles containing densities corresponding to RECQL5 from both datasets were combined, reverted to original particles and 3D-refined (Extended Data Fig. 3k). Following signal subtraction with a soft mask on PAF and focused 3D-classification without alignment (Extended Data Fig. 3l), particles containing PAF densities were selected, reverted to original particles and imported into CryoSPARC for local refinement. This resulted in the final EC*- RECQL5 reconstruction with 314,016 particles at an overall resolution of 2.0 Å (Extended Data Fig. 3m).

To improve the resolution of elongation factors, soft masks were applied individually onto SPT6^Core^, SPT6^Core^-Pol II^Stalk^, SPT6^tSH2^, PAF (CTR9-SKI8 region), PAF1-LEO1, SPT4 and SPT5 (KOW2-KOW3 region). Following particle subtraction, 3D classification without alignment was performed for SPT6^tSH2^, PAF1-LEO1, SPT4 and SPT5 (KOW2-KOW3 region), resulting in medium resolution maps of each region. These maps allowed docking of existing structures and AlphaFold^64^ predicted models of the elongation factors into respective regions. For SPT6^Core^, SPT6^Core^-Pol II^Stalk^ and PAF (CTR9-SKI8 region), 3D classification did not find any classes lacking densities corresponding to the elongation factors. Therefore, all subtracted particles were imported into CryoSPARC for local refinement, resulting in a resolution at 2.5 Å for all three maps of SPT6^core^, SPT6^Core^-Pol II^Stalk^ and PAF (CTR9-SKI8 region) (Extended Data Fig. 3n-p).

For the dataset of the EC-TCR-RECQL5 complex, particles after cleanup in 2D classification were CTF-refined (Extended Data Fig. 10a, b), followed by focused 3D- classification without alignment using a soft mask on the RECQL5 IRI domain to separate EC-TCR-RECQL5 particles from EC-TCR only particles (Extended Data Fig. 10c). Particles containing densities of RECQL5 were selected and 3D-refined (Extended Data Fig. 10d).

Subsequent focused 3D-classification without alignment using a soft mask on TCR were performed to exclude EC-RECQL5 only particles (Extended Data Fig. 10e). This results in the final EC-TCR-RECQL5 reconstruction with 19,458 particles at an overall resolution of 3.5 Å (Extended Data Fig. 10f).

Local resolution of the maps was estimated using CryoSPARC. All resolution calculations were based on the gold-standard Fourier Shell Correlation (FSC) using the FSC = 0.143 criterion. A summary of all EM reconstructions obtained in this paper is listed in Extended Data Table 1.

### Model building and refinement

Initial models of *S. scrofa* Pol II (PDB: 7B0Y^35^) and EC* (PDB: 6GMH^13^) as well as AlphaFold^64^ predictions of RECQL5 and elongation factors were rigid-body fitted into the overall map in Chimera^65^. The Pol II and RECQL5 models were manually adjusted in Coot^66^ using the overall map of EC*-RECQL5 (2.0 Å) and the structure was real-space refined in PHENIX^67^. The SPT6^core^ model was manually adjusted in Coot using the local refined SPT6 map (2.5 Å) and real-space refined in PHENIX. The refined SPT6^core^ model was then fitted into the local refined SPT6^core^-Pol II^stalk^ map (2.5 Å), adjusted in Coot and real-space refined in PHENIX. The PAF model was manually adjusted in Coot using local refined PAF map (2.5 Å) and the structure was real-space refined in PHENIX. The rest of the elongation factors were rigid-body docked into the overall map in Chimera. The resulting complete model of EC*-RECQL5 was then real-space refined in the overall map using PHENIX and structure restraints of Pol II, SPT6^core^-Pol II^stalk^ and PAF.

An additional density was found to bind the tSH2 domain of SPT6, and the tSH2 domain is in a horizontal conformation compared to previous EC* structures^13^ (Extended Data Fig. 3q). De novo model building using ModelAngelo^68^ identified this unassigned density to be CDC73. This allowed us to identify a previously uncharacterized interface between the tSH2 domain of SPT6 and CDC73 of PAF (Extended Data Fig. 5e), consistent with biochemical analysis in yeast^69^.

The EC-RECQL5 model was fitted into the overall map of EC-RECQL5, manually adjusted in Coot and real-space refined in PHENIX.

The EC-RECQL5 model from EC*-RECQL5 and the TCR model (8B3D)^38^ were rigid-body fitted into the overall map of EC-TCR-RECQL5 and real-space refined using adp refinement in PHENIX. Figures were generated using PyMOL (The PyMOL Molecular Graphics System, Version 2.0 Schrödinger, LLC.) and Chimera X^70^. Sequence alignment was performed using Jalview^71^.

### Crosslinking-coupled mass spectrometry

Samples for crosslinking-coupled mass spectrometry were prepared essentially as described for cryo-EM studies of EC*-RECQL5, except that peak fractions of EC* were combined and incubated with 1.5x molar excess of RECQL5 on ice for 30 min, crosslinked with 1 mM BS3 on ice for 1 hour and quenched with 50 mM Tris-HCl pH 8.0.

The quenched solution was reduced with 5 mM DTT and alkylated with 20 mM idoacetamide. SP3 protocol as described in^72,73^ was used to clean up and buffer exchange the reaction. Briefly, the complex was washed with ethanol, resuspended in 100 mM NH_4_HCO_3_ and digested with trypsin (Promega) at an enzyme-to-substrate ratio of 1:20, and 0.1% ProteaseMAX (Promega) overnight at 37 °C. Digested peptides were purified using HyperSep SpinTip P-20 C18 columns (ThermoScientific), eluted with 60% (v/v) acetonitrile (ACN), and dried using Speed Vac Plus (Savant). Dried peptides were resuspended in 30% (v/v) ACN and separated using a Superdex 30 Increase 3.2/300 column (Cytiva) at a flow rate of 10 μL/min in 30% (v/v) ACN and 0.1% (v/v) trifluoroacetic acid. Fractions containing crosslinked peptides were collected and dried with Speed Vac Plus (Savant). Dried peptides were suspended in 3% (v/v) ACN and 0.1 % (v/v) formic acid and analysed by nanoscale capillary LC-MS/MS using an Ultimate U3000 HPLC (ThermoScientific) to deliver a flow of 300 nl/min. Peptides were trapped on a C18 Acclaim PepMap100, 5 μm, 0.3 μm x 5 mm cartridge (ThermoScientific) before separation on Aurora Ultimate C18, 1.7 μm, 75 μm x 25 cm (Ionopticks). Peptides were eluted on a 90-min gradient and interfaced via an EasySpray ionisation source to a QExactive Plus mass spectrometer (ThermoScientific). Data were acquired in data-dependent mode using a Top-15 method, where high resolution full mass scans were carried out (R = 70,000, m/z 400 – 1500) followed by higher energy collision dissociation (HCD) of 30 V. The tandem mass spectra were recorded (R = 60,000, automatic gain control target = 5e5, maximum injection time = 100 ms, isolation window = 1.2 m/z, dynamic exclusion = 40 s).

Xcalibur raw files were converted to MGF files using ProteoWizard^74^ and crosslinks were analysed by XiSearch^75^. Search conditions used 3 maximum missed cleavages with a minimum peptide length of 5. Variable modifications used were carbmidomethylation of cysteine (57.02146 Da) and oxidation of methionine (15.99491 Da). False discovery rate was set to 5%. The sequence database was assembled from all proteins within the complex. Crosslink sites were visualized with xiVIEW^76^ and PyXlinkViewer plugin^77^ in PyMOL Version 2.0.

## Data availability

The cryo-EM reconstructions and final models were deposited with the EMDB under accession code EMD-52440 and the PDB under accession code 9HVO for the EC-RECQL5 complex, EMD-52443 (overall), EMD-52441 (SPT6-stalk), EMD-52442 (PAF) and PDB 9HVQ for the EC*-RECQL5 complex, and EMD-52449 and PDB 9HWG for the EC-TCR-RECQL5 complex.

## Acknowledgments

We thank all members of the Zhang lab for discussion, particularly G.A. Arroyo for participating in the initial purification of RECQL5. We thank P. Cramer for the tagged Pol II human cell line. We thank the LMB EM facility and scientific computing for maintaining the microscopes and the computing cluster, respectively and K. Turton for maintaining the insect cell facility. We thank S. Aibara for suggestions on model building and manuscript, and D. Barford, L. Passmore and M. Hegde for suggestions on the manuscript. This work was funded by the Medical Research Council, as part of United Kingdom Research and Innovation (also known as UK Research and Innovation) with the MRC file reference number MC_UP_1201/30 for S.Z. For open access, the MRC Laboratory of Molecular Biology has applied a CC BY public copyright license to any Author Accepted Manuscript version arising.

## Author contributions

L.Z. and S.Z. designed and performed experiments, collected and analyzed cryo-EM data. Y.G. performed experiments. T.M. and C.F. performed mass spectrometry. A.T.V. and S.Z. initiated the project. S.Z. supervised the project and built the models. S.Z. and L.Z. wrote the manuscript with input from all other authors.

## Competing interests

Authors declare that they have no competing interests.

**Correspondence and request of materials** should be addressed to S.Z. under a material transfer agreement with the MRC Laboratory of Molecular Biology.

---

**Extended Data Fig. 1.**
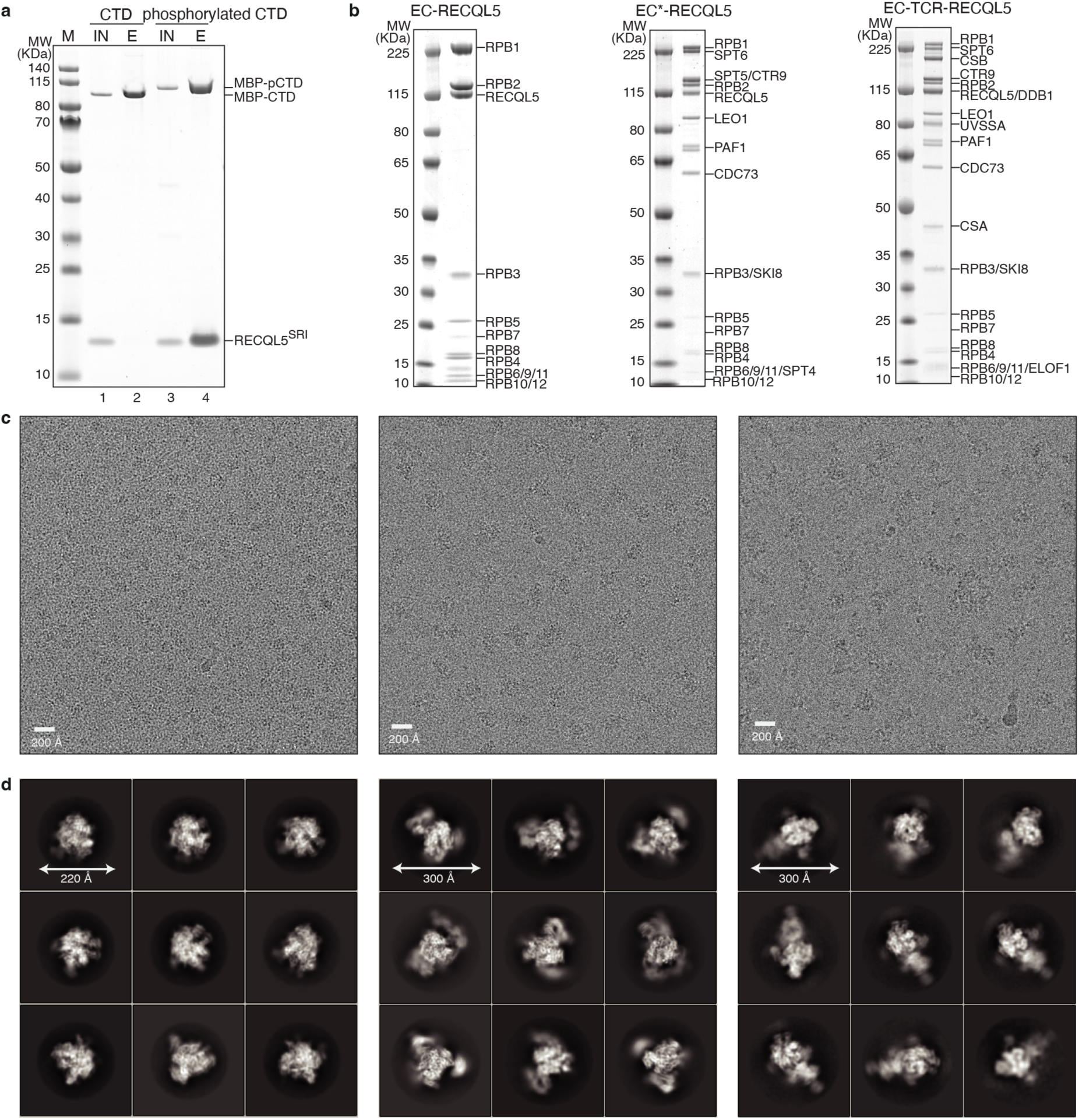
Preparation of the EC-RECQL5, EC*-RECQL5 and EC-TCR-RECQL5 complexes. **a**, Pulldown of the SRI domain of RECQL5 with MBP-tagged Pol II CTD. Phosphorylation of the CTD (pCTD) is required for its interaction with the SRI domain. **b**, Assembled EC-RECQL5 (left), EC*-RECQL5 (middle), and EC-TCR-RECQL5 (right) complexes on 4-12% NuPAGE Bis-Tris gels run in MOPS, stained with Instant Blue. **c**, Representative micrographs of EC-RECQL5 (left), EC*-RECQL5 (middle) and EC-TCR-RECQL5 (right) complexes collected on the 300 kV Titan Krios with a Falcon4i detector in electron counting mode. **d**, Two-dimensional averages of the EC-RECQL5 (left), EC*-RECQL5 (middle) and EC-TCR-RECQL5 (right) complexes.

**Extended Data Fig. 2.**
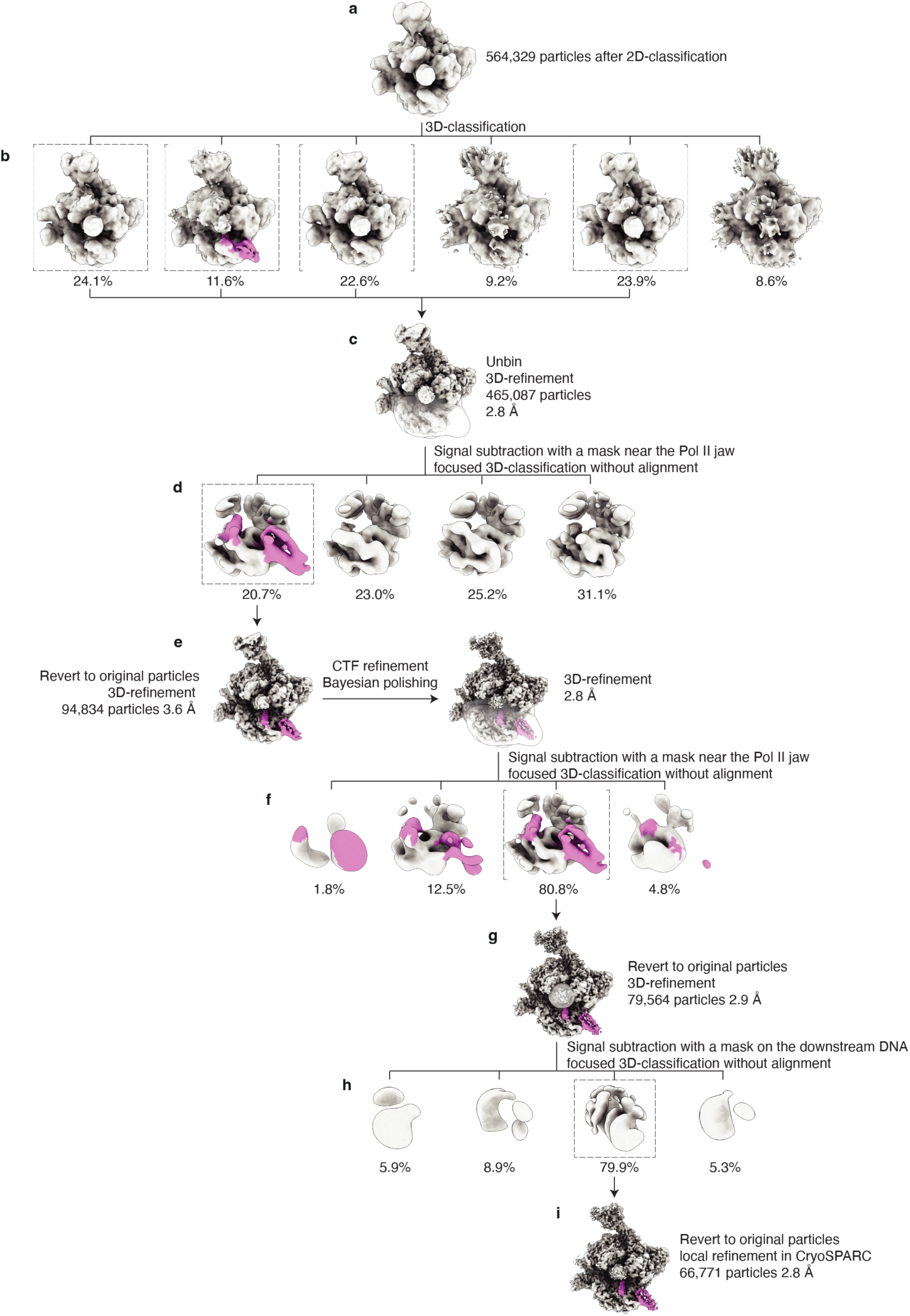
Cryo-EM data processing of the EC-RECQL5 complex. After removing junk particles with 2D-classification, the remaining particles were 3D-refined and further cleaned up using local 3D-classification (**a**, **b**). Good particles were extracted without binning and 3D-refined (**c**). Following signal subtraction with a soft mask near the Pol II jaw and focused 3D-classification without alignment (**d**), particles containing densities of RECQL5 were reverted to original particles and 3D-refined before CTF refinement and Bayesian polishing (**e**). A second round of signal subtraction and focused 3D-classification on RECQL5 were performed to further remove the EC-only particles (**f**, **g**). Subsequent signal subtraction with a soft mask on the downstream DNA and focused 3D-classification were performed to exclude the apo Pol II-RECQL5 particles (**h**). This resulted in a final EC-RECQL5 reconstruction at 2.8 Å resolution (**i**). The cryo-EM densities were coloured according to Fig. 1d.

**Extended Data Fig. 3.**
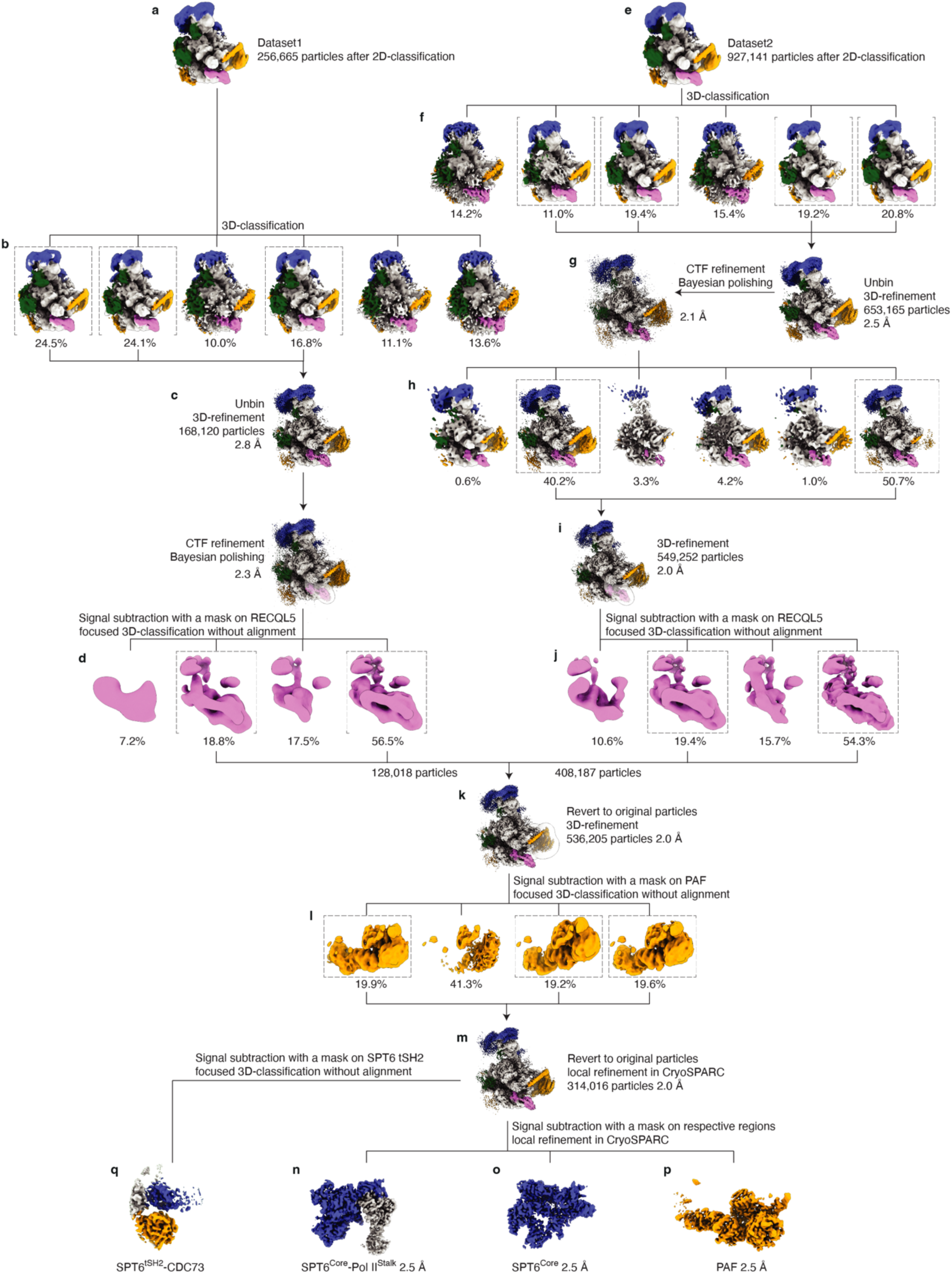
Cryo-EM data processing of the EC*-RECQL5 complex. 2D-classification, followed by 3D-refinement and local 3D-classification were performed to remove bad particles from the datasets (**a**, **b**, **e**, **f**). Remaining particles underwent CTF refinement and Bayesian polishing separately (**c**, **g**). Polished particles in Dataset 2 were subjected to an additional round of local 3D-classification to exclude junk particles (**h**, **i**). Following signal subtraction of Pol II and focused 3D-classification on RECQL5 (**d**, **j**), particles containing densities of RECQL5 from both datasets were combined, reverted to original particles and 3D-refined (**k**). This is followed by signal subtraction and focused 3D-classification with a soft mask on PAF (**l**). Particles containing densities of PAF were reverted to original particles and 3D-refined, resulting in a final EC*-RECQL5 reconstruction at an overall resolution of 2.0 Å (**m**). Particle subtraction followed by local refinement improved the local resolution of SPT6^core^, SPT6^core^-Pol II^stalk^ and PAF (CTR9-SKI8 region) (**n**-**p**). Signal subtraction and focused 3D-classification without alignment on SPT6^tSH2^ identified a previously unassigned density of CDC73 bound to the tSH2 domain of SPT6 (**q**). The cryo-EM densities were coloured according to Fig. 1e.

**Extended Data Fig 4.**
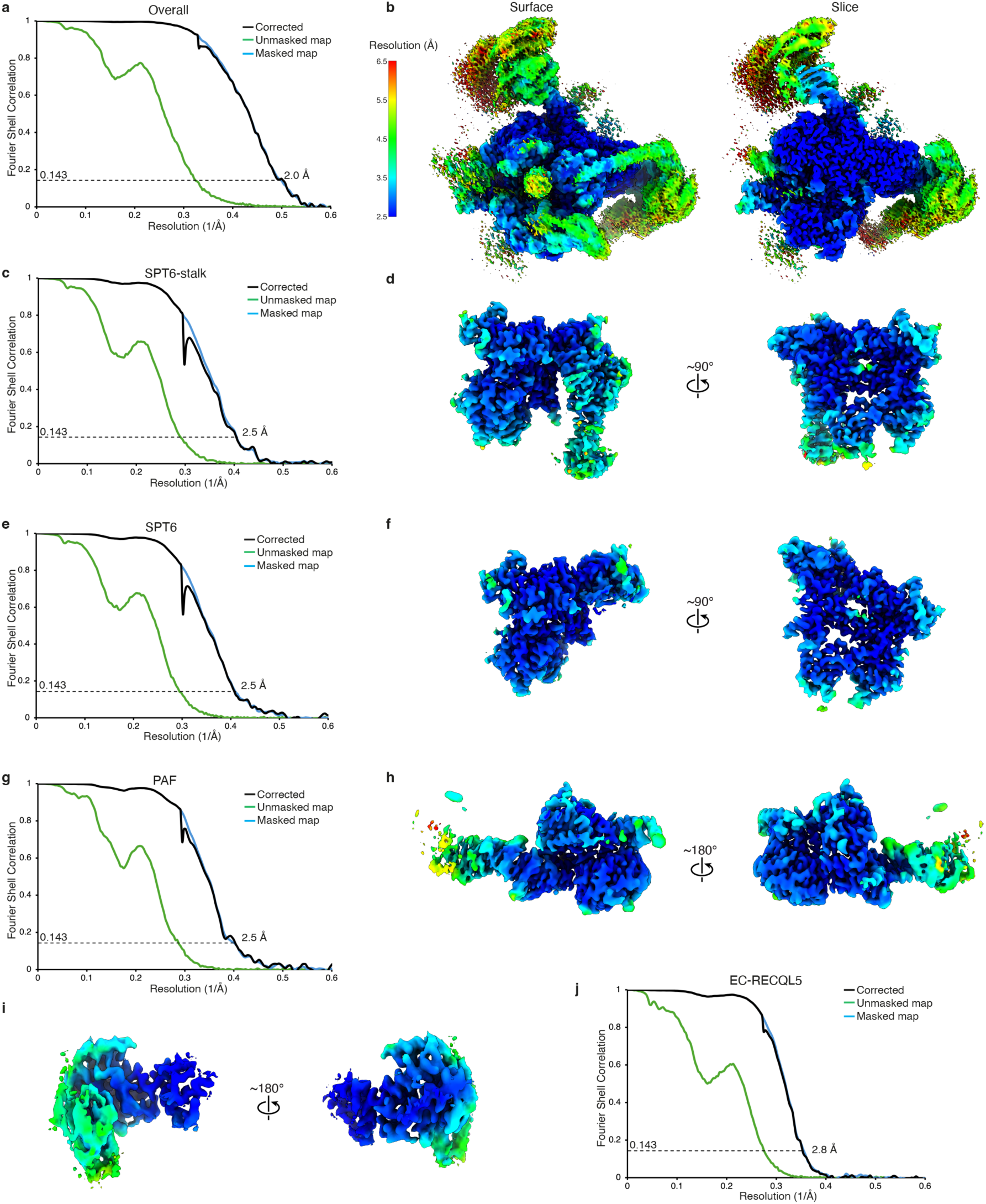
Local resolution estimation and FSC curves of EC-RECQL5 and EC*-RECQL5 maps. **a**, Gold-standard Fourier Shell Correlation (FSC) curves of the overall map of EC*-RECQL5. **b**, Local resolution estimations of the overall map in both surface and slice views showed that the Pol II core is beyond 2.5 Å and RECQL5 is around 3.5 Å. **c**-**h**, FSC curves of the focused refined maps of SPT6^Core^-Pol II^Stalk^ (**c**), SPT6^Core^ (**e**) and PAF (**g**). Local resolution estimation of the focused refined maps showed improved density of SPT6^Core^-Pol II^Stalk^ (**d**), SPT6^Core^ (**f**) and PAF (**h**) to mostly beyond 2.5 Å. **i**, Local resolution estimation of the RECQL5-Pol II interface. The brake helix interface with the Pol II jaw is resolved to 3 Å, revealing bulky side chain densities. The KIX domain is more mobile, at around 4 Å. **j**, FSC curves of the overall map of EC-RECQL5.

**Extended Data Fig 5.**
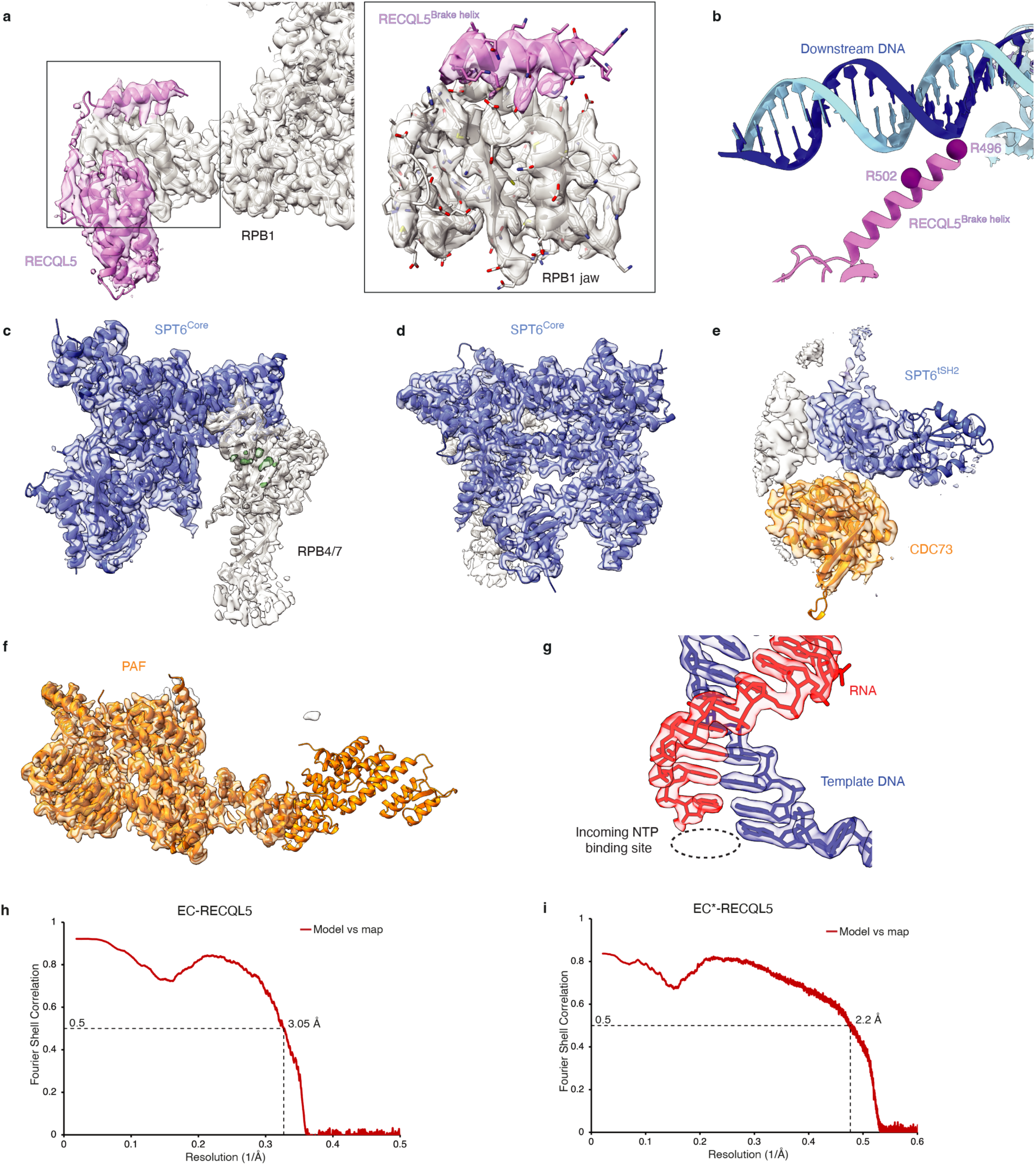
Cryo-EM densities of the EC*-RECQL5 reconstruction. **a**, Cryo-EM densities of the overall map showing the interaction between the RECQL5 IRI domain and the RPB1 jaw domain. The views and colours are the same as in Fig. 2A. **b**, Close-up view of RECQL5 brake helix (pink) protruding towards the downstream DNA (blue and cyan). Cα atoms of R496 and R502 residues are shown in magenta spheres. **c**, **d**, **f**, Cryo-EM densities of the local refined maps with subtracted particles for SPT6^Core^-Pol II^Stalk^ (**c**), SPT6^Core^ (**d**) and PAF (**f**). **e**, Cryo-EM densities showing the SPT6^tSH2^-CDC73 interaction. **g**, Cryo-EM densities at the active centre reveal the post-translocated state that can accept incoming nucleotide. **h**, **i**, Model versus map FSC for the overall map of EC-RECQL5 (**h**) and EC*-RECQL5 (**i**) using the FSC standard of 0.5. The cryo-EM densities were coloured according to Fig. 1e.

**Extended Data Fig 6.**
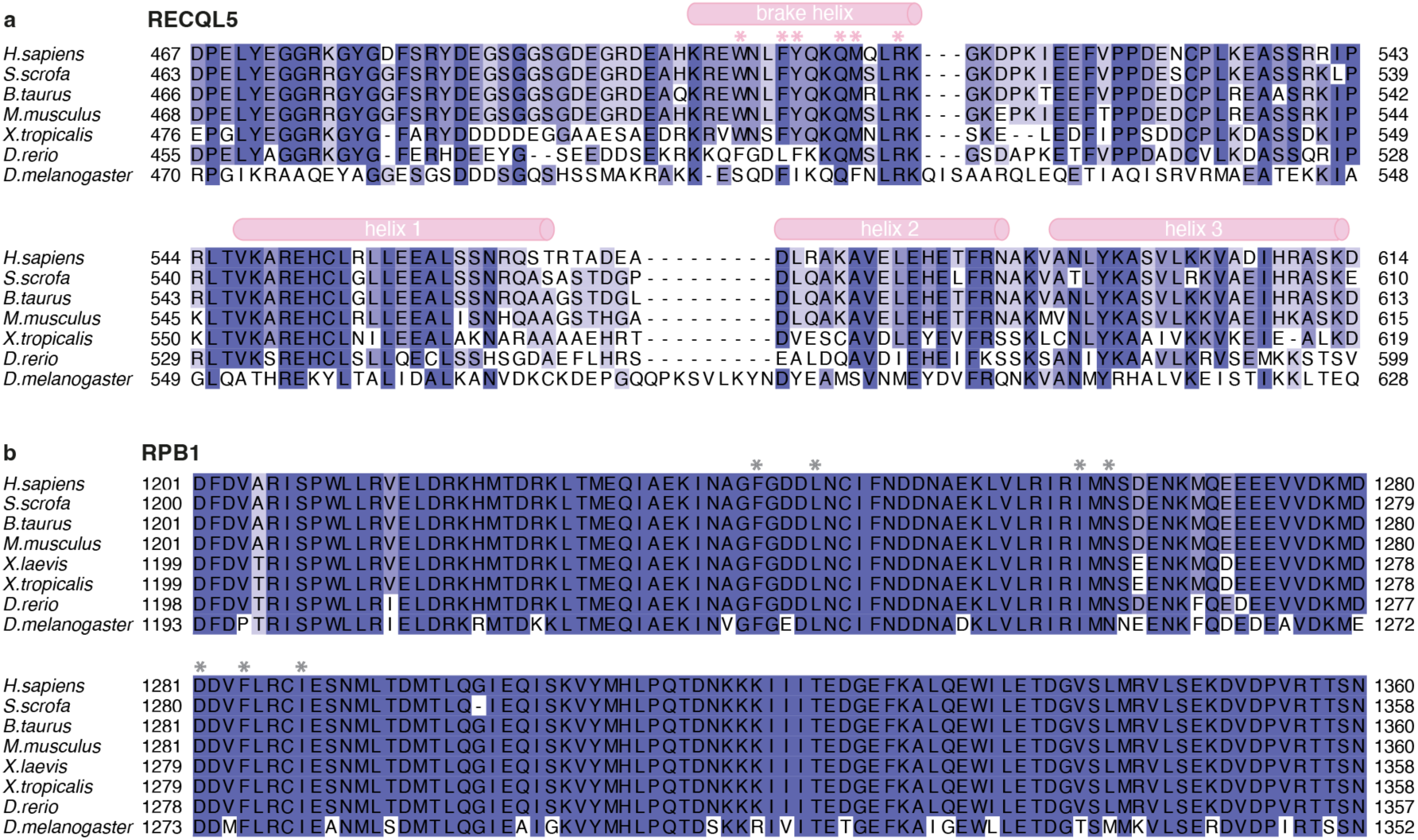
Sequence alignments of RECQL5 and RPB1 at the interface region. **a**, Sequence alignment of the RECQL5 IRI domain, with the position of the brake helix and the three α-helices of the KIX domain shown as cylinders. Residues of the brake helix in contact with RPB1 are marked with pink stars. **b**, Sequence alignment of the RPB1 jaw domain. Residues that contact the RECQL5 brake helix are marked with grey stars.

**Extended Data Fig. 7.**
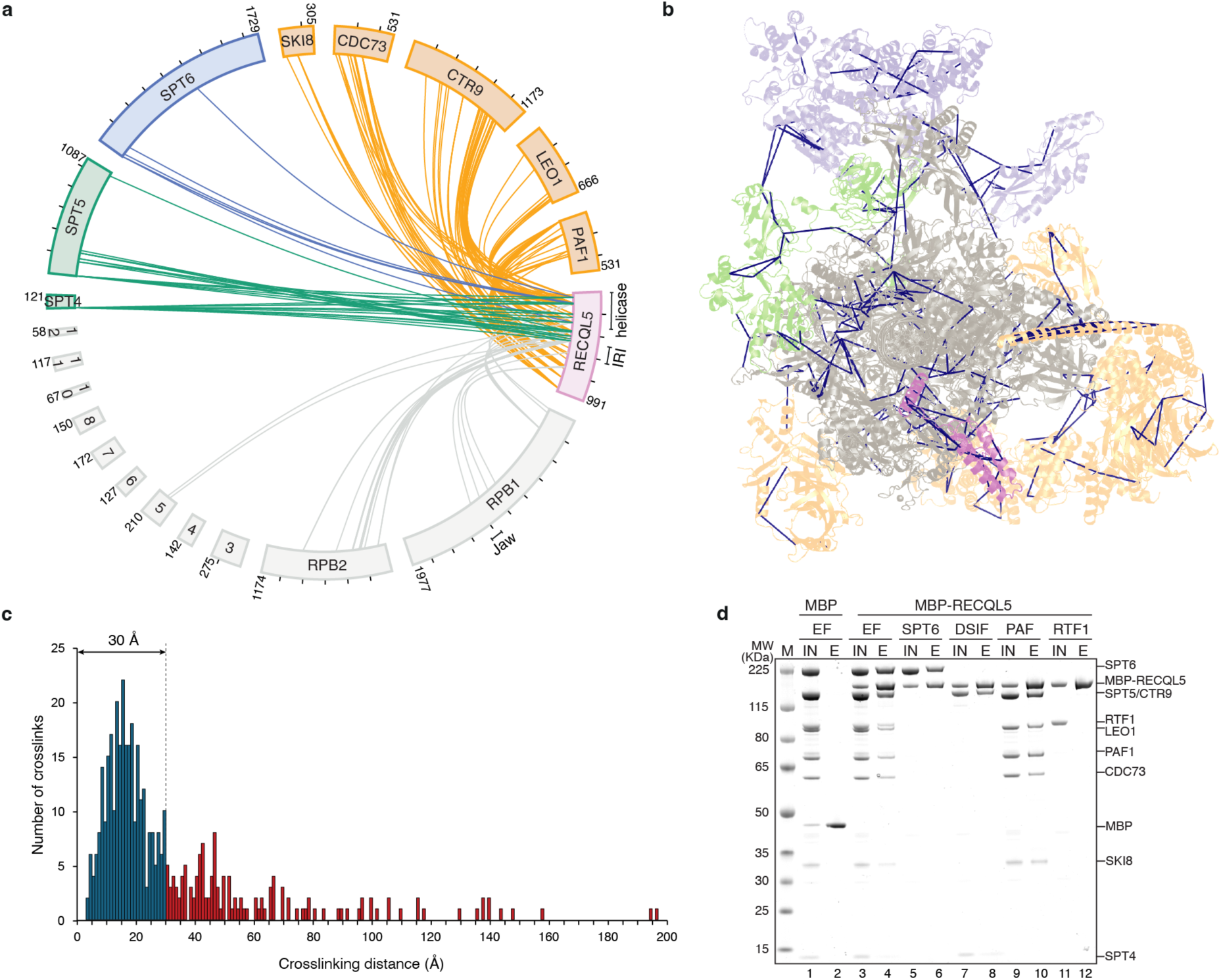
Crosslinking-coupled mass spectrometry analysis of the EC*-RECQL5 complex. **a**, Crosslinking network of the EC*-RECQL5 complex with BS3. Crosslinks of RECQL5 (pink) with Pol II (grey) and elongation factors SPT6 (blue), DSIF (green) and PAF (orange) are shown. Interlinks were detected between RECQL5 and RPB1, RPB2, RPB5, PAF components, and the N-termini of SPT4, SPT5 and SPT6. **b**, Crosslinks mapped onto the EC*-RECQL5 structure. Crosslinks among all subunits of the EC*-RECQL5 complex within the permitted distance of 30 Å are shown. **c**, Histogram of the number of crosslinks and distances between crosslinked Cα pairs mapped onto the EC*-RECQL5 structure. 69% of mapped crosslinks fall within the permitted crosslinking distance of 30 Å (blue). Crosslinks outside the permitted distance (red) are likely due to complex aggregation and flexibility. **d**, Pulldown of RECQL5 with elongation factors (EF) SPT6, DSIF, PAF and RTF1 using an MBP tag on RECQL5. SPT6 formed a stoichiometric complex with RECQL5, while DSIF and PAF bound to RECQL5 at reduced levels. RTF1, a loosely associated factor of PAF, showed no interaction with RECQL5. IN: input, E: elution. Pulldown was repeated in triplicate.

**Extended Data Fig. 8.**
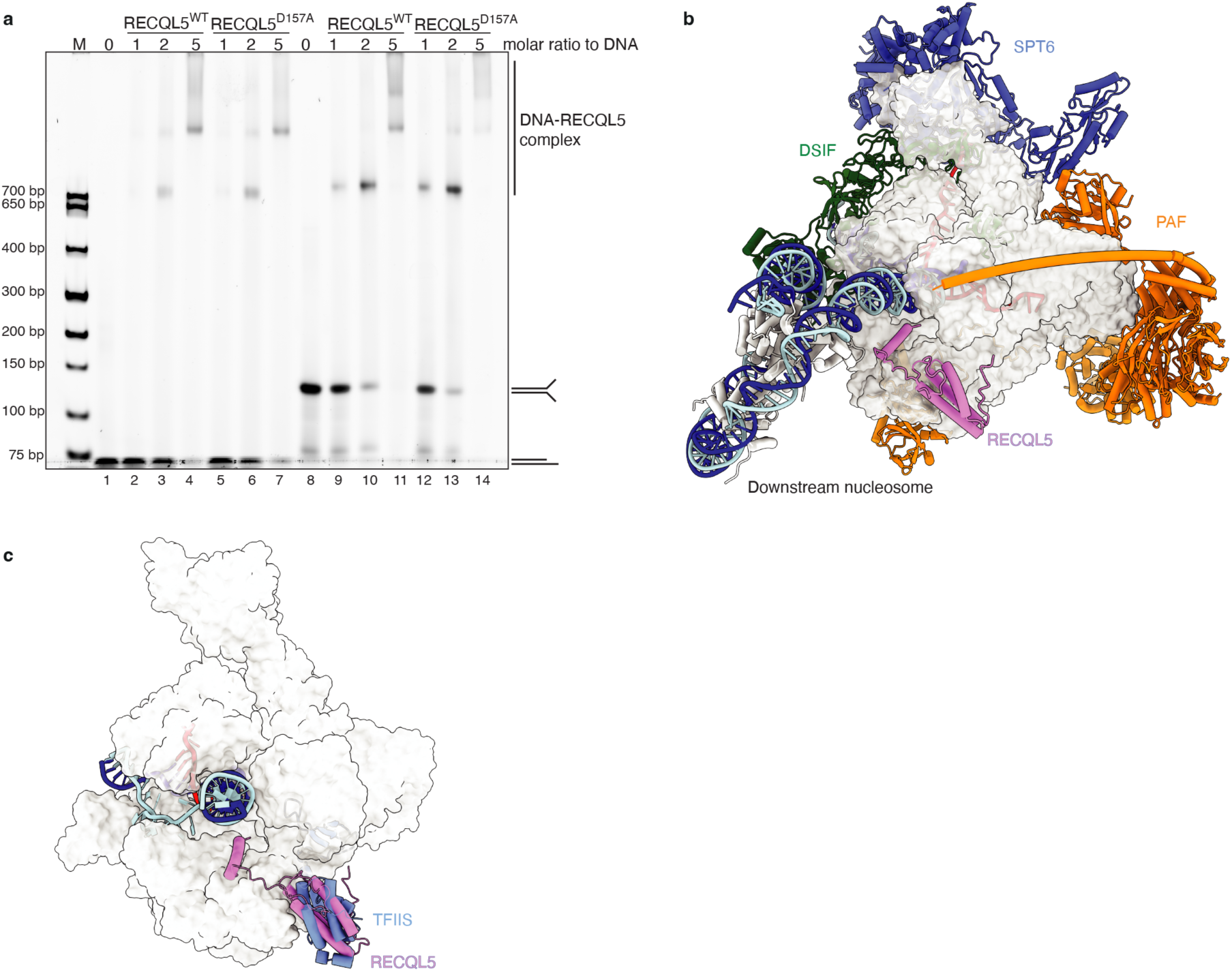
Superposition of the EC-RECQL5 structure with transcription elongation complexes. **a**, Electromobility shift assay (EMSA) of wildtype and D157A mutant RECQL5. D157A mutation does not affect RECQL5 binding to the DNA. The assay was repeated in triplicate. **b**, Superposition of the EC*-RECQL5 complex with EC*-nucleosome (PDB: 8A3Y)^36^ on Pol II showed no clash between RECQL5 and the downstream nucleosome. EC* and RECQL5 are depicted as described in Fig. 1e and his tones are shown in white. **c**, Superposition of the EC-RECQL5 complex with EC-TFIIS (PDB: 8A40)^37^ on Pol II showed that the RECQL5 KIX domain and TFIIS share overlapping binding site on the Pol II jaw. TFIIS is shown in light blue.

**Extended Data Fig. 9.**
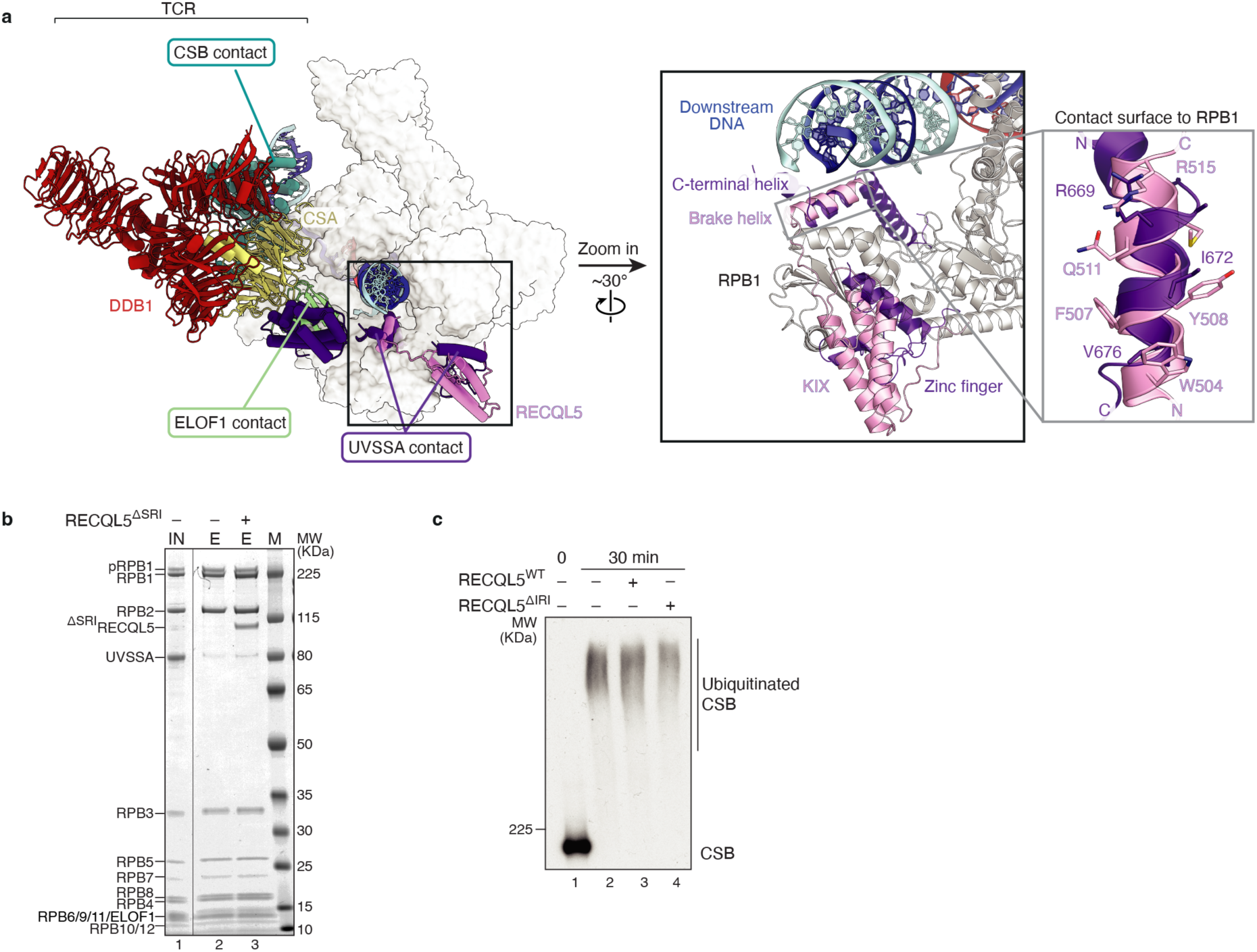
RECQL5 and UVSSA compete for the same binding site on Pol II. **a**, Superposition of the EC-RECQL5 structure with the EC-TCR structure (PDB: 8B3D)^38^ reveals that the brake helix of RECQL5 (pink) and the C-terminal hinged helix of UVSSA (dark purple) bind to the same site on the Pol II jaw, while the RECQL5 KIX domain and the UVSSA zinc finger domain occupy the same binding site. The TCR complex engages the Pol II additionally with CSB and ELOF1. EC-TCR complex and RECQL5 are depicted as in Fig. 3d. Right panel: The RECQL5 brake helix (pink) and the UVSSA hinged helix (dark purple) have opposite directionality. **b**, Competition assay of RECQL5^ΔSRI^ and UVSSA for Pol II binding. The assembled transcription complex EC was pre-incubated with UVSSA before addition of RECQL5^ΔSRI^. RECQL5 has a stronger interaction with Pol II compared to UVSSA. Pulldown was performed in duplicate. **c**, CSB ubiquitination by CRL4 is not affected by the presence of either wildtype or ΔIRI mutant RECQL5.

**Extended Data Fig 10.**
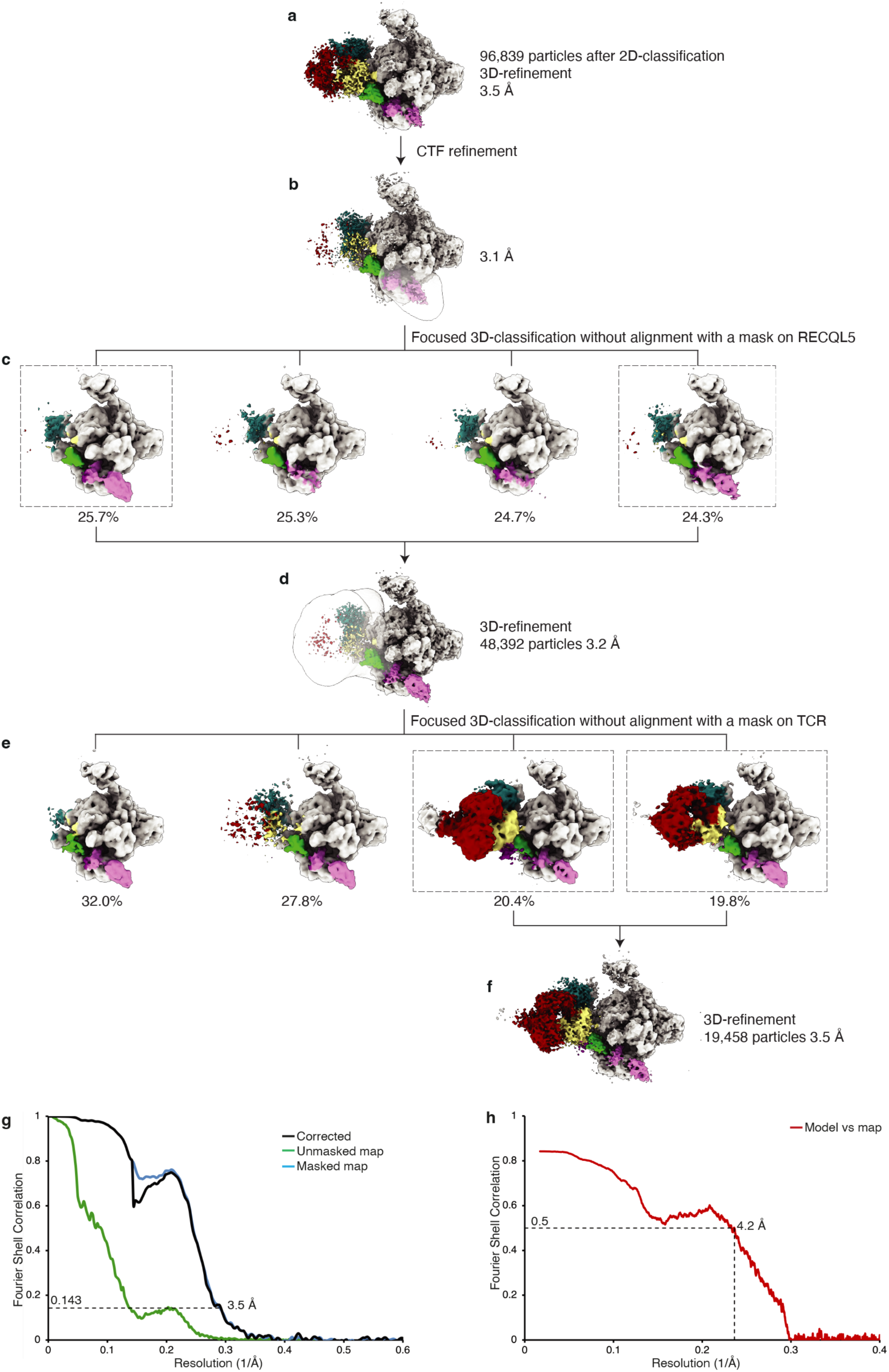
Cryo-EM data processing of the EC-TCR-RECQL5 complex. After removing bad particles with 2D-classification, the remaining particles were CTF-refined (**a**, **b**). Focused 3D-classification without alignment were performed using a soft mask on RECQL5 (**c**). Particles containing densities of RECQL5 were combined and 3D-refined (**d**). Following focused 3D-classification on TCR were performed to remove the EC-RECQL5 only particles (**e)**, resulting in the final EC-TCR-RECQL5 reconstruction at 3.5 Å resolution (**f**). (**g**) FSC curves of the final map for the EC-TCR-RECQL5 complex. (**h**) Model versus map FSC for the overall map of EC-TCR-RECQL5 using the FSC standard of 0.5. The cryo-EM densities were coloured according to Fig. 3d.

**Extended Data Table 1.** Statistics of cryo-EM reconstructions and structural models.

|  | EC*-RECQL5 |  |  | EC-RECQL5 | EC-TCR-RECQL5 |
| --- | --- | --- | --- | --- | --- |
|  | Overall<br>(EMD-52443) | SPT6-stalk<br>(EMD-52441) | PAF<br>(EMD-52442) | Overall<br>(EMD-52440) | Overall<br>(EMD-52449) |
| **Data collection and processing** |  |  |  |  |  |
| Microscope | Titan Krios |  |  | Titan Krios | Titan Krios |
| Voltage (kV) | 300 |  |  | 300 | 300 |
| Camera | Falcon 4i |  |  | Falcon 4i | Falcon 4i |
| Magnification | 96 000 x |  |  | 96 000 x | 96 000 x |
| Pixel size (Å/pixel) | 0.8156 |  |  | 0.815 | 0.8156 |
| Electron exposure (e<sup>-</sup>/Å<sup>2</sup>) | 39.9 |  |  | 42.7 | 36.6 |
| Exposure rate (e<sup>-</sup>/Å<sup>2</sup>/frame) | 0.997 |  |  | 1.06 | 0.915 |
| Number of frames per movie | 40 |  |  | 40 | 40 |
| Defocus range (µm) | 0.5-2.0 |  |  | 0.5-2.0 | 0.5-2.0 |
| Automation software | EPU |  |  | EPU | EPU |
| Symmetry imposed | C1 |  |  | C1 | C1 |
| Initial particle numbers | 1,183,806 |  |  | 564,329 | 96,839 |
| Final particle numbers | 314,016 |  |  | 66,771 | 19,458 |
| Map sharpening *B* factor (Å<sup>2</sup>) | -35.9 | -68.1 | -67 | -84.7 | -51.9 |
| Map resolution (Å, FSC=0.143) | 2.0 | 2.5 | 2.5 | 2.8 | 3.5 |
| **Refinement** |  |  |  |  |  |
| Initial models used (PDB) | 6TED, 7B0Y, AlphaFold models |  |  | 7B0Y, AlphaFold | 9HVQ, 8B3D |
| Model resolution (Å) | 2.2 |  |  | 3.05 | 4.2 |
| Model composition |  |  |  |  |  |
| Non-hydrogen atoms | 63,185 |  |  | 33,905 | 51248 |
| Protein residues | 7,623 |  |  | 4,014 | 6,203 |
| Nucleic acid residues | 86 |  |  | 86 | 92 |
| Ligands | Zn:8 Mg:1 |  |  | Zn:8 Mg:1 | Zn:10 Mg:2 |
| *B* factors (Å<sup>2</sup>) |  |  |  |  |  |
| Protein | 230.67 |  |  | 27.26 | 221.89 |
| Nucleotide | 171.29 |  |  | 87.11 | 239.57 |
| Ligand | 83.58 |  |  | 81.41 | 204.87 |
| R.m.s. deviations |  |  |  |  |  |
| Bond lengths (Å) | 0.003 |  |  | 0.004 | 0.009 |
| Bond angles (°) | 0.529 |  |  | 0.497 | 0.817 |
| Validation |  |  |  |  |  |
| MolProbity score | 1.39 |  |  | 1.45 | 1.76 |
| Clash score | 5.29 |  |  | 4.64 | 7.56 |
| Poor rotamers (%) | 0.62 |  |  | 0 | 1.28 |
| CB deviations (%) | 0 |  |  | 0 | 0 |
| Ramachandran plot |  |  |  |  |  |
| Favored (%) | 97.49 |  |  | 96.63 | 96.15 |
| Allowed (%) | 2.44 |  |  | 3.37 | 3.83 |
| Outliers (%) | 0.07 |  |  | 0.0 | 0.02 |
| PDB code | 9HVQ |  |  | 9HVO | 9HWG |

**Extended Data Table 2.**
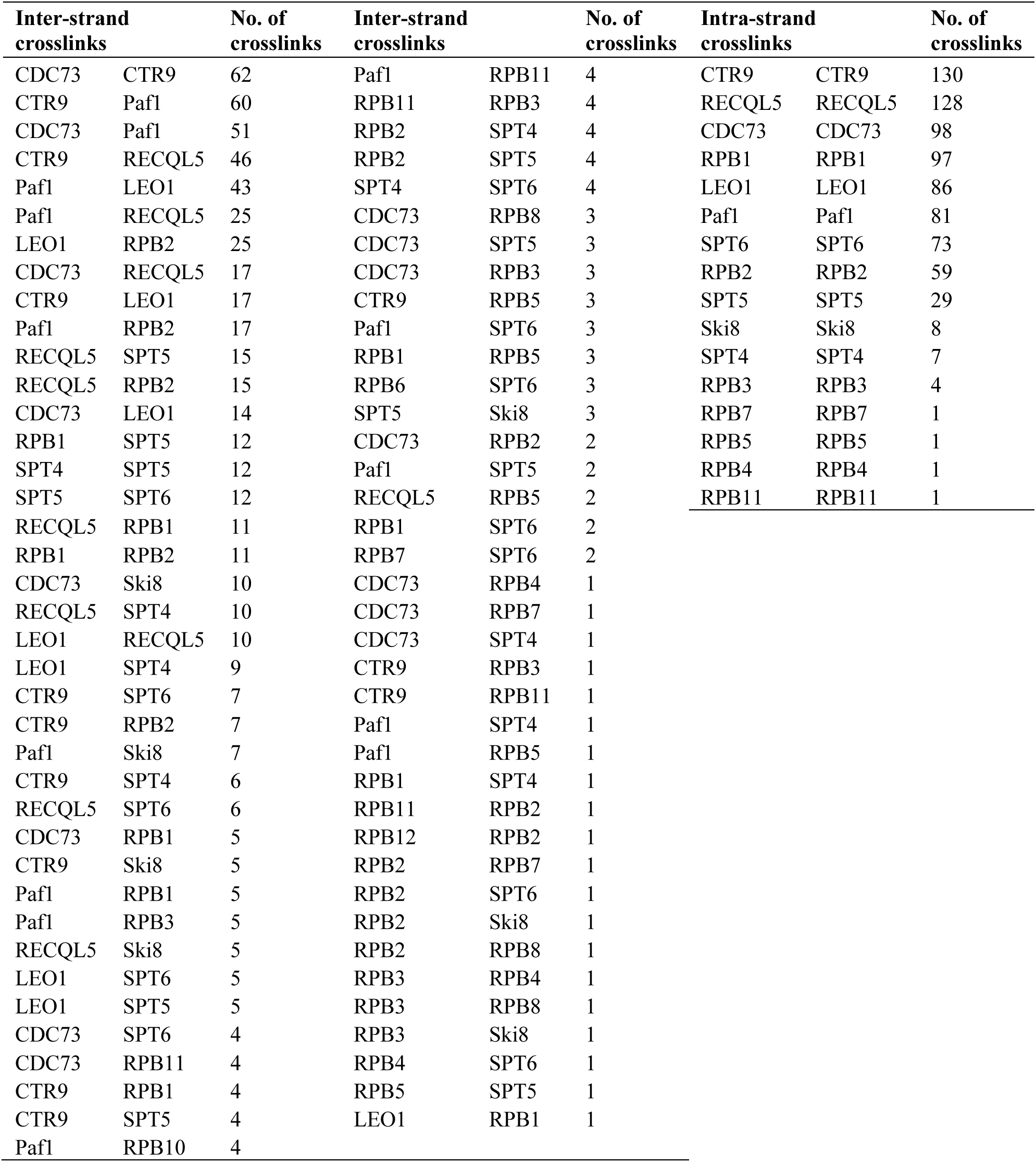
Crosslinking mass-spectrometry of the EC*-RECQL5 complex.

